# Pharmacologic and Genetic Modulation of PERK-associated Stress Signaling Enhances Regulatory T Cell Responses and Attenuates Intestinal Inflammation

**DOI:** 10.64898/2026.11.20.764961

**Authors:** Kibrom M. Alula, Tom T. Nguyen, Natarajan Balasubramaniyan, Tzu Phang, Tian Liu, Julian G. Contreras, Gabriela R. Kobak, Paul Jedlicka, Edwin F. de Zoeten

## Abstract

**Background and aims:** The unfolded protein response (UPR) and integrated stress response (ISR) coordinate cellular adaptation to stress, with PERK-eIF2α-ATF4 linking ER stress to ISR signaling. We investigated how PERK-associated stress signaling regulates regulatory T cell (Treg) differentiation, phenotype, and function.

**Materials and methods:** Murine CD4^+^ T cells were exposed to ER-stress inducers and pharmacologic modulators of PERK/ISR-associated signaling. CD4^+^ T cells from WT-CD4^cre^ and PERK-CD4^cre^ mice were differentiated under Treg-polarizing conditions and analyzed by flow cytometry. Treg suppressive activity was assessed by effector T-cell proliferation assays. Bulk and single-cell RNA sequencing defined stress-associated transcriptional programs and PERK-associated Treg heterogeneity. Systemic CCT020312 (CCT) treatment was evaluated in TNF^ΔARE+/−^ mice.

**Results:** Global UPR activation increased FoxP3^+^ Treg differentiation and enriched Treg-associated transcriptional programs and PERK-associated signatures. PERK-deficient cells showed reduced FoxP3^+^ Treg frequencies under vehicle, tunicamycin, and CCT conditions and consistently reduced Neuropilin-1 expression, whereas CTLA-4 and TIGIT were increased, indicating selective phenotypic remodeling. Pharmacologic PERK/ISR modulation altered FoxP3 protein abundance and Treg-mediated control of effector T-cell proliferation. Single-cell RNA sequencing identified a PERK-high intestinal Treg state enriched for stress-adaptive programs. In vivo, systemic CCT increased lamina propria FoxP3^+^ and IL-10^+^ Treg responses and was associated with reduced ileal inflammation.

**Conclusions:** Pharmacologic and genetic findings support a role for CD4^+^ T-cell PERK signaling in shaping stress-dependent Treg differentiation and phenotype. Differential regulation of FoxP3/Neuropilin-1 versus CTLA-4/TIGIT suggests that PERK selectively influences distinct components of the Treg program during intestinal inflammation.

## 1 Introduction

The endoplasmic reticulum (ER) is central to cellular homeostasis, coordinating protein folding, maturation, lipid synthesis, and calcium storage (1). Perturbations in ER function, such as increased secretory demand, metabolic stress, or inflammatory signaling, lead to accumulation of unfolded or misfolded proteins and activation of the unfolded protein response (UPR) (2). The UPR is an evolutionarily conserved ER stress-sensing network composed of three principal branches: inositol- requiring enzyme 1 (IRE1), protein kinase RNA-like ER kinase (PERK), and activating transcription factor 6 (ATF6) (3). Together, these pathways restore proteostasis by transiently attenuating protein translation, enhancing expression of molecular chaperones, and promoting degradation of misfolded proteins. While initially characterized in highly secretory cells such as pancreatic β-cells and plasma cells (4), UPR signaling is now recognized as a critical regulator of immune cell function (5, 6).

The integrated stress response (ISR), by contrast, is a broader adaptive signaling program that responds to multiple forms of cellular stress, including amino acid deprivation, viral infection, oxidative stress, and ER stress (7–9), through phosphorylation of eukaryotic initiation factor 2α (eIF2α) and selective translation of stress-responsive transcripts such as ATF4 (8, 10). PERK is therefore unique in that it functions as both a canonical branch of the UPR and a kinase that links ER stress to ISR activation (9). Upon ER stress, PERK phosphorylates eIF2α, transiently reducing global protein synthesis while selectively enhancing translation of ATF4 and downstream adaptive targets (11, 12). This PERK-eIF2α-ATF4 axis promotes amino acid biosynthesis, redox balance, and proteostatic resilience. In innate immune cells, PERK signaling modulates macrophage survival and inflammatory polarization (13) and influences dendritic cell activation (14). In T cells, PERK activity has been shown to fine-tune activation thresholds and metabolic programming (15). However, the role of PERK signaling in regulatory T cells (Tregs) remains incompletely defined.

Tregs are essential for maintaining immune tolerance and preventing excessive inflammation (16, 17). These cells operate in metabolically demanding and highly inflammatory environments, particularly at barrier sites such as the intestinal mucosa (18, 19). Within these settings, Tregs must sustain expression of lineage-defining factors such as FoxP3 and immunosuppressive mediators including IL-10 and CTLA-4, while simultaneously adapting to cytokine-driven and metabolic stress. FoxP3 stability is tightly regulated at transcriptional, epigenetic, and post-translational levels, and disruption of these regulatory layers can lead to loss of suppressive identity and acquisition of effector-like phenotypes (20, 21). ER homeostasis is therefore likely critical for preserving Treg stability and function (22, 23). Indeed, chronic or unresolved ER stress has been associated with reduced FoxP3 expression and impaired suppressive capacity (24). Xu et al. demonstrated that inflammatory cytokines induce ER stress in Tregs, destabilizing FoxP3, whereas the E3 ligase Hrd1 maintains Treg identity by restraining excessive UPR activation (22). Conversely, activation of certain UPR branches, especially the IRE1α/XBP1 axis, has been reported to enhance Treg fitness in inflamed tissues (25). These observations suggest that UPR signaling can either support or undermine Treg biology depending on context, intensity, and pathway balance.

Emerging evidence now implicates the PERK arm specifically in shaping T cell responses. Feng et al. reported that PERK signaling enhances T cell receptor-induced activation in Tregs (26), while studies in other T cell subsets have linked PERK-dependent translational control to metabolic adaptation and cytokine production (13, 27). Notably, ER stress induction via thapsigargin (Thaps) was shown to increase IL-10 transcription in human T cell clones through PERK-eIF2α signaling, and this effect was modulated by manipulation of eIF2α phosphorylation (28). These findings raise the possibility that PERK activation may reinforce Treg suppressive programs by enabling adaptive stress responses rather than driving maladaptive ER stress (29).

Despite these insights, the therapeutic potential of selectively targeting PERK to bolster Treg function has not been systematically examined. This question is particularly relevant in inflammatory bowel disease (IBD), where Treg dysfunction contributes to chronic mucosal inflammation (30). This is partly due to disruption of the Th17/Treg balance, to which microbiota-derived signals are important contributors (31). In addition, Tregs residing in inflamed intestinal tissue exhibit evidence of heightened ER stress, suggesting that their capacity to maintain proteostasis may be overwhelmed (22). We therefore hypothesized that controlled activation of the PERK pathway could enhance Treg lineage stability and suppressive function by promoting adaptive eIF2α-ATF4 stress signaling without triggering maladaptive CHOP-dominated responses.

In this study, we investigated the contribution of PERK-associated stress signaling to Treg differentiation, phenotype, and function using complementary pharmacologic and genetic approaches. General UPR inducers and pharmacologic modulators of PERK/ISR-associated signaling were combined with CD4-specific PERK deletion to determine the extent to which Treg responses depend on PERK. Bulk and single-cell RNA-sequencing analyses were used to define stress- associated transcriptional programs and PERK-related heterogeneity within the Treg compartment. Finally, systemic CCT treatment was evaluated in TNF^ΔARE+/-^ mice to determine whether pharmacologic stress-pathway modulation is associated with altered intestinal Treg responses and inflammation. Together, these approaches identify PERK as a context-dependent contributor to Treg stress adaptation while revealing differential regulation of distinct components of the Treg phenotype.

## 2 Materials and methods

### 2.1 Mice

Nine- to thirteen-week-old male C57BL/6 mice were purchased from The Jackson Laboratory and maintained under specific pathogen-free conditions. Male TNF^ΔARE /^ mice, 16-18 weeks of age, were maintained under the same conditions and used for in vivo studies. For genetic assessment of CD4^+^ T-cell PERK signaling, PERK-floxed mice were crossed with CD4-Cre mice to generate PERK-CD4^cre^ mice and corresponding WT-CD4^cre^ controls, 16-18 weeks of age male mice. Genotypes were confirmed by PCR using Cre universal and PERKflox primer sets, with the expected genotype-specific PCR products shown in Supplementary Figure 1. Fecal samples were confirmed negative for Helicobacter species, protozoa, and helminths. All animal procedures were approved by the Institutional Animal Care and Use Committee (IACUC) of the University of Colorado Anschutz Medical Campus and conducted in accordance with institutional guidelines.

### 2.2 Cell isolation and UPR modulation

Spleen and mesenteric lymph nodes (MLN) were extracted from euthanized mice as previously reported (32). Briefly, single-cell suspension of splenocytes and MLN were isolated through a 70 µm cell strainer. Splenic red blood cells were lysed by 3-minute exposure to ammonium chloride lysing reagent (ACK Lysis Buffer; Invitrogen, Carlsbad, CA). CD4^+^ cells were isolated according to manufacturer’s instructions (EasySepTM Mouse CD4^+^ T cell Isolation Kit, 19852, STEMCELL Technologies, British Columbia, Canada). Isolated CD4^+^ cells were further separated into CD25^+^ (Tregs) and CD25^neg^ (non-Tregs) cells according the manufacturer’s instructions (EastSepTM Mouse CD25 Regulatory T cell Positive Selection Kit, 18782, STEMCELL Technologies, British Columbia, Canada). Non-Treg cells were counted with a cell counter (Cellometer Auto T4, Nexcelom, Bioscience, CA, USA). 1 x 10^6^ cells were plated per well and incubated with the following UPR pathway modulators for 4 hrs at 37°C: 1X PBS Vehicle control (Veh), 0.15 ng/mL of Tunicamycin (Tun, UPR inducer; sc-3506, SantaCruz, CA, USA), 0.5 g/L low-glucose (Glu, UPR inducer; 15023- 021, Gibco, NY, USA), 50 nM Thapsigargin (Thaps, UPR inducer; T7458, ThermoFisher, OR, USA), 800 nM of CCT020312 (CCT, pharmacologic PERK-pathway activator; 324879, Millipore Sigma, CA, USA), 1 µM of GSK2606414 (GSK, pharmacologic PERK kinase inhibitor; 516535, Millipore Sigma, CA, USA), or 1µg/mL Salubrinal (Sal, inhibitor of eIF2α dephosphorylation; SML0951, Millipore Sigma, CA, USA) (32). Because these compounds differ in target selectivity and can influence signaling beyond PERK itself, pharmacologic treatment was interpreted as modulation of PERK-associated/ISR signaling rather than definitive evidence of PERK-specific activity unless otherwise supported by pathway measurements. For genetic-validation experiments, CD4^+^CD25^neg^ cells isolated from WT-CD4^cre^ and PERK-CD4^cre^ mice were exposed to the same Veh, Tun, CCT, GSK, and Sal treatment conditions for 4 hrs followed by Treg conversion for 72 hrs.

### 2.3 Treg conversion assay

The CD4^+^CD25^neg^ cells were converted into CD4^+^CD25^+^FoxP3^+^ cells by incubating them in complete media (RPMI 1640 + L-glutamine, 10-040-CV, Corning Inc., Corning NY, USA) containing 10 % FBS (MT35011CV, Corning, VA, USA), 100 IU Penicillin (15140122, Gibco, NY, USA), and 100 µg/ml Streptomycin (15140-122, Gibco, NY, USA), IL-2 (5 IU/mL; 212-12, PEPROTECH, NJ, USA), anti-CD28 (1 µg/mL; 102102, BioLegend, CA, USA), TGFβ1 (2 ng/mL; 100-21C, PEPROTECH, NJ, USA), and plate-bound anti-CD3 (1 µg/mL; 100302, BioLegend, CA, USA) for 72 hrs (18). Cells were lysed for Western Blot (immunoblot). WT-CD4^cre^ and PERK- CD4^cre^ CD4^+^CD25^neg^ cells were subjected to identical Treg-polarizing conditions to permit direct genotype-by-treatment comparisons.

### 2.4 Flow cytometry

The remaining cells were subdivided into two: Cells were isolated from the spleen and MLN as in section 2.2 and the converted CD4^+^CD25^+^ (section 2.3) and the original CD4^+^CD25^neg^ cells were assessed by Flow Cytometry for FoxP3 expression by flow cytometry (Cytek Aurora R0027, Fremont, CA, USA) after staining with FoxP3 APC (17-5773-82, Invitrogen, CA, USA), TruStain FcX (Fc block, 101320; BioLegend, CA, USA), CD4 PerCP/Cy5.5 (100434, BioLegend, CA, USA), AmCyan Live/Dead Aqua Fluorescent reactive dye (34966A, BioLegend, CA, USA), Neuropilin-1 PE/Cy7 (145212, BioLegend, CA, USA), TIGIT PE Cy7 (142108, BioLegend, CA, USA), ICOS APC Cy7 (313530, BioLegend, CA, USA), PD-1 FITC (135214, BioLegend, CA, USA), and CTLA- 4 PE (106306, BioLegend, CA, USA). Flow-cytometric analysis was performed using FlowJo software (Tree Star Inc., Ashland, OR, USA). Viable singlets were first gated on CD4 and CD8 expression. Tregs were defined as CD4^+^FoxP3^+^ cells, and expression of Neuropilin-1, ICOS, CTLA- 4, and TIGIT was subsequently quantified within the CD4^+^FoxP3^+^ Treg population. For analysis of effector CD4^+^ T-cell cytokine responses in ileal LP samples, CD4^+^FoxP3^neg^ cells were evaluated for intracellular IFNγ and IL-17 expression. Representative gating strategies for CD8^+^ T cells and FoxP3^neg^ effector CD4^+^ T-cell cytokine populations are shown in Supplementary Figure 2. The same hierarchical gating strategy was applied across treatment and genotype groups. A complete list of antibodies, reagents, kits, instruments, and software used in this study is provided in Supplementary Table 1.

### 2.5 Bulk RNA-sequencing

Bulk RNA-seq was performed on isolated CD4 T cells obtained from the spleen and MLN of mice and treated ex vivo with either Tun or Veh for 4 hours. Following treatment, total RNA was extracted using the RNeasy Mini Kit (74104, QIAGEN, MD, USA). Subsequent sample processing was performed at the Genomics Shared Resources Core, University of Colorado Anschutz Medical Campus. RNA purity, concentration, and integrity were assessed using a NanoDrop One spectrophotometer (ND-ONE-W, Thermo Scientific, WI, USA) and an Agilent TapeStation 4200 system (G2991BA, Agilent Technologies, CA, USA). For library construction, 200 ng of total RNA was used as input for the Universal Plus™ mRNA-Seq Library Preparation Kit with NuQuant (0520, TECAN, Männedorf, Switzerland). Libraries were sequenced on a NovaSeq X Plus platform (Illumina, CA, USA) to generate 150 bp paired-end reads, with a target depth of 40 million paired- end reads per sample. Raw sequencing data were de-multiplexed using bcl2fastq. Bulk RNA-seq data analysis was performed beginning with FASTQ files for each sample. Transcript quantification was carried out using Salmon, and data were imported into R with tximport. Sample-level quality control was assessed using DESeq2. Normalization was performed by library size adjustment followed by low-expression filtering. Principal component analysis (PCA) was applied for dimensionality reduction. Differential expression analysis was conducted with DESeq2, and pathway enrichment was performed using clusterProfiler and fgsea. Results were visualized with PCA plots, volcano plots, and heatmaps.

### 2.6 Single cell RNA-sequencing

Single-cell RNA-seq was performed on live LP cells isolated from the ileal small bowel of TNF^ΔARE+/-^ mice. Ileal tissue was harvested, opened longitudinally, washed to remove luminal contents, and cut into small pieces. Epithelial cells were removed by incubation in dissociation buffer, after which the remaining tissue was enzymatically digested to isolate LP leukocytes and generate a single-cell suspension. Viable cells were enriched by flow cytometric sorting using live/dead discrimination with 7AAD staining (420404, BioLegend, CA, USA). Single-cell libraries were generated using the 10x Genomics Chromium platform according to the manufacturer’s protocol (Single Cell 3′ v3.1 chemistry). Briefly, cells were loaded onto a Chromium Controller to generate gel bead-in-emulsion (GEM) droplets, followed by reverse transcription and library construction. Libraries were sequenced on an Illumina platform to achieve a target depth of approximately 50,000-100,000 reads per cell. Raw sequencing data were processed using Cell Ranger (10x Genomics) for alignment to the mouse reference genome (mm10), barcode assignment, and gene counting. Downstream analysis was performed using Seurat (v4). Cells with fewer than 200 or more than 2,500 detected features and cells with >5% mitochondrial reads were excluded. Suspected doublets were removed prior to downstream analysis. Data were normalized using the SCTransform workflow or log-normalization as appropriate. Principal-component analysis was performed and the first 10 PCs were used for dimensionality reduction and clustering at a resolution of 0.2, followed by UMAP visualization. Tregs were identified based on canonical marker expression (e.g., *Foxp3*, *Il2ra*), and PERK module scores were calculated using curated gene sets (*Eif2ak3*, *Atf4*, *Ddit3*, *Ppp1r15a*, etc.) via the AddModuleScore function. Differential expression between PERK- high and PERK-low Tregs was assessed using the Wilcoxon rank-sum test with false discovery rate correction. Heatmaps and violin plots were generated to visualize gene expression patterns.

### 2.7 Suppression assay

Treg in vitro suppression assay was performed as previously described (32). Briefly, antigen- presenting cells (APC) were isolated from WT C57BL/6 mouse using the mouse CD90.2 positive selection kit II (18951, STEMCELL, British Columbia, Canada) per manufacturer’s instruction. Then, the isolated APC cells were irradiated with 30 Gray (MultiRad350, PRECISION X-ray Irradiation, CT, USA). 50,000 CD4^+^CD25^neg^ effector T cells were stained with Violet-labeled CellTrace (C34557, Thermo Fisher, MA, USA) and stimulated with anti-CD3 (1 µg/mL; 100302, BioLegend, CA, USA) at varying effector-to-Treg ratios (1:1 to 1:32). Each well also contained 75,000 irradiated syngeneic antigen-presenting cells (MultiRad350, PRECISION X-ray Irradiation, CT, USA). After 72 hours of incubation at 37°C, the suppression of effector cell proliferation was assessed by staining cells for CD4^+^ T cells with CD4 PerCP/Cy5.5 (100434, BioLegend, CA, USA) and defining the proliferation peaks based on CellTrace dilution peaks by flow cytometry.

### 2.8 Immunoblot analysis

Proteins from the converted Treg samples (section 2.3) were extracted with NP40 Cell Lysis buffer (FNN0021, Invitrogen, MA, USA) containing 10% of HaltTM Protease & Phosphatase Single-Use Inhibitor cocktail (78442, Thermo Scientific, IL, USA). PierceTM BCA Protein Assay Kit (23225, Thermo Scientific, IL, USA) was used to determine protein concentration for each sample per manufacturer’s instructions. Proteins were then separated by SDS-PAGE and transferred onto PVDF membranes using the Trans-Blot Turbo Transfer system (17001915, Bio-Rad, CA, USA). Membranes were blocked in 5% BSA in Tris-buffered saline with 0.1% Tween-20 (TBS-T) for 1 hour at room temperature. The membranes were then incubated overnight at 4°C with the following primary antibodies: Phospho-PERK (pPERK; 1:1000, MA5-15033, Invitrogen, MA, USA), Phospho- eIF2α (1:1000, D9G8, Cell Signaling, MA, USA), FoxP3 (0.5 µg/mL, 126402, BioLegend, CA, USA), Il-10 (1:1000, ARC9102, Invitrogen, MA, USA), ATF4 (1 µg/mL; PA5-68801, Invitrogen, MA, USA), CHOP (1:2000, 15204-1-AP, Proteintech, IL, USA), and GAPDH (loading control; 1:2000, PA1-987, Invitrogen, MA, USA). After washing with TBS-T, membranes were incubated with goat anti-rabbit secondary antibody conjugated to horseradish peroxidase (HRP; 1:10,000, A27036, Invitrogen, MA, USA) and HRP Goat anti-rat IgG (1:5000, 405405, BioLegend, CA, USA) for 1 hour at room temperature. Protein bands were visualized using enhanced chemiluminescence (ECL) detection reagents (170-5060, Bio-Rad, CA, USA) according to the manufacturer’s instructions and imaged using the ChemiDocTM MP Imaging System (Bio-Rad, CA, USA).

### 2.9 In vivo CCT treatment in TNF^ΔARE+/-^ mice

Male TNF^ΔARE /^ mice (16-18 weeks of age) were maintained under specific pathogen-free conditions and randomly assigned to receive either the CCT or vehicle control. CCT was dissolved in DMSO and diluted in sterile 1× PBS prior to administration, and mice were treated by intraperitoneal (i.p.) injection at a dose of 4 mg/kg daily for 14 consecutive days. Control animals received an equivalent volume of DMSO diluted in 1× PBS. The experimental timeline is illustrated in Figure 7A. At the end of the treatment period, mice were euthanized and the ileum was harvested. Leukocytes from LP were isolated following removal of epithelial cells and mechanical dissociation of the tissue, followed by enzymatic digestion to generate single-cell suspensions. Cells were stained for flow cytometric analysis of CD4 T cells, regulatory T cells (CD4 FoxP3 ), and intracellular IL-10 expression using standard fixation and permeabilization protocols. Data were acquired on a flow cytometer and analyzed using FlowJo software. In parallel, sections of distal ileum from each mouse were fixed in 10% neutral-buffered formalin, paraffin-embedded, sectioned, and stained with hematoxylin and eosin (H&E). Histologic assessment was performed by a gastrointestinal pathologist (Co Author P.J.) blinded to treatment assignment. Ileal inflammation was evaluated using a previously described semi-quantitative scoring system (32, 33). Accordingly, three histologic parameters were assessed: active inflammation, defined by granulocytes/neutrophil infiltration; chronic inflammation, defined by lymphocyte, plasma cell, and macrophage infiltration within the mucosa and submucosa; and villus distortion, defined by flattening and/or widening of the normal villous architecture. Each parameter was scored from 0 (normal) to 3 (severe), and the individual component scores were summed to generate a total ileal inflammation score ranging from 0 to 9. Representative images were acquired for analysis.

### 2.10 Statistical analysis

Statistical analyses were performed using GraphPad Prism (version 11.1.0, MA, USA) and R (version 4.6.1). For two-group comparisons, unpaired two-tailed Student’s t-tests were used for normally distributed data were applied. For genetic-validation experiments comparing WT-CD4^cre^ and PERK-CD4^cre^ cells across pharmacologic conditions, two-way ANOVA was performed with genotype and treatment as factors, followed by Sidak’s multiple-comparisons test. Genotype x treatment interactions were evaluated to determine whether pharmacologic responses differed according to PERK genotype. For other multiple-group comparisons, one-way or two-way ANOVA with the indicated post hoc test was used as appropriate. Data are presented as mean ± SEM, and p < 0.05 was considered statistically significant.

A summary of the experimental design, treatment conditions, primary readouts, and biological replication for each major experiment is provided in Supplementary Table 2.

## 3 Results

### 3.1 Broad UPR activation enhances FoxP3 Treg differentiation

We utilized a Treg conversion assay to determine whether ER stress influences regulatory T cell differentiation. For this assessment we used CD4 T cells exposed to multiple pharmacologic and metabolic pan-UPR activators. Tun-induced ER stress through inhibition of N-linked glycosylation, significantly increased the proportion of CD4 FoxP3 Tregs compared with Veh (Figure 1A). Similarly, metabolic ER stress induced by low-glucose conditions (0.5 g/L) modestly but significantly enhanced Treg frequency (Figure 1B). Perturbation of ER calcium homeostasis using Thaps also resulted in a marked increase in FoxP3 Tregs (Figure 1C). Across all treatments, total CD4 T cell frequencies remained unchanged, indicating that ER stress selectively promotes Treg differentiation rather than global CD4 T cell expansion. Because all three global ER stress activators activate multiple arms of the UPR, these findings indicate that the tested ER stress conditions increased the proportion of FoxP3^+^ cells generated under Treg-polarizing conditions.

**FIGURE 1.**
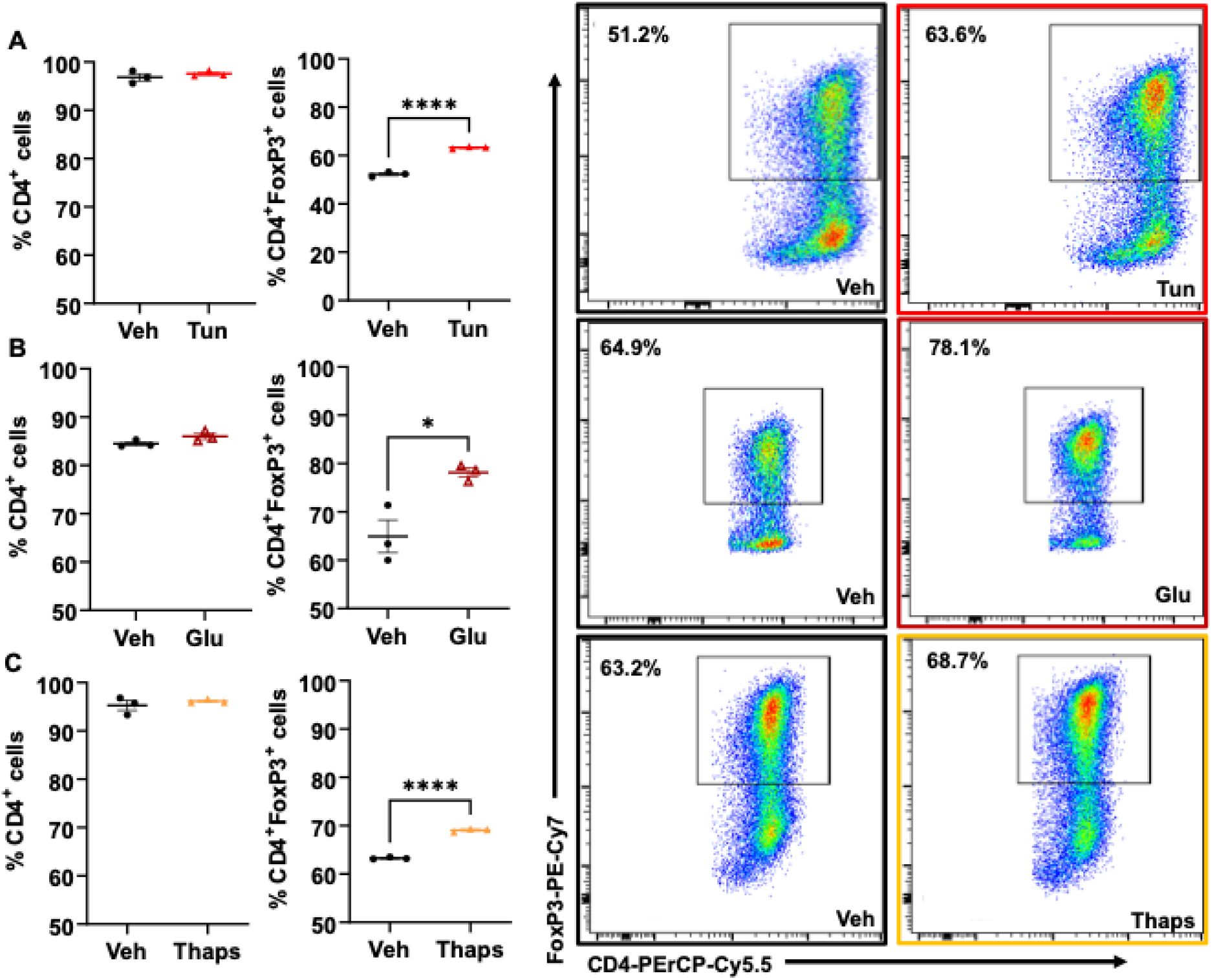
Broad UPR activation increases FoxP3^+^ Treg differentiation. **(A)** Quantification of total CD4^+^ cells and CD4^+^FoxP3^+^ Tregs following treatment with vehicle (Veh) or tunicamycin (Tun), with representative flow-cytometry plots shown at right. **(B)** Quantification of total CD4^+^ cells and CD4^+^FoxP3^+^ Tregs following treatment with Veh or low-glucose conditions (Glu), with representative flow-cytometry plots shown at right. **(C)** Quantification of total CD4^+^ cells and CD4^+^FoxP3^+^ Tregs following treatment with Veh or thapsigargin (Thaps), with representative flow- cytometry plots shown at right. Each data point represents an independent biological replicate, and data are presented as mean ± SEM. Statistical significance was determined using an unpaired two- tailed Student’s *t*-test. \**p* < 0.05; \*\*\*\**p* < 0.0001.

### 3.2 Tunicamycin-induced ER stress enriches Treg transcriptional programs and PERK- associated pathway activity

To define transcriptional changes associated with ER stress-enhanced Treg differentiation, bulk RNA sequencing was performed on WT CD4 T cells treated with Tun or Veh for 4 hrs at 37^0^C. Principal component analysis demonstrated clear segregation of Tun-treated and Veh-treated samples, indicating global transcriptional reprogramming (Figure 2A). Differential expression analysis revealed significant upregulation of multiple Treg-associated genes and coordinated repression of distinct gene subsets (Figure 2B). Heatmap analysis confirmed increased expression of canonical Treg lineage and functional markers, including *Il2ra (CD25)*, *Foxp3*, *Ikzf2 (Helios)*, *Il10*, *Pdcd1*, *Tnfrsf4*, and *Tnfrsf18* (Figure 2C). Functional pathway analysis demonstrated enrichment of programs governing Treg differentiation and suppressive function, with concurrent reduction of pathways associated with proliferation and tissue adaptation (Figure 2D). Importantly, gene set variation analysis (GSVA) revealed enrichment of a PERK-associated transcriptional signature in Tun-treated samples (Figure 2E). Moreover, PERK-associated module scores positively correlated with Treg transcriptional enrichment in Tun-treated cells (Figure 2F), demonstrating an association between enrichment of PERK-related stress response signatures and reinforcement of the Treg transcriptional program.

**FIGURE 2.**
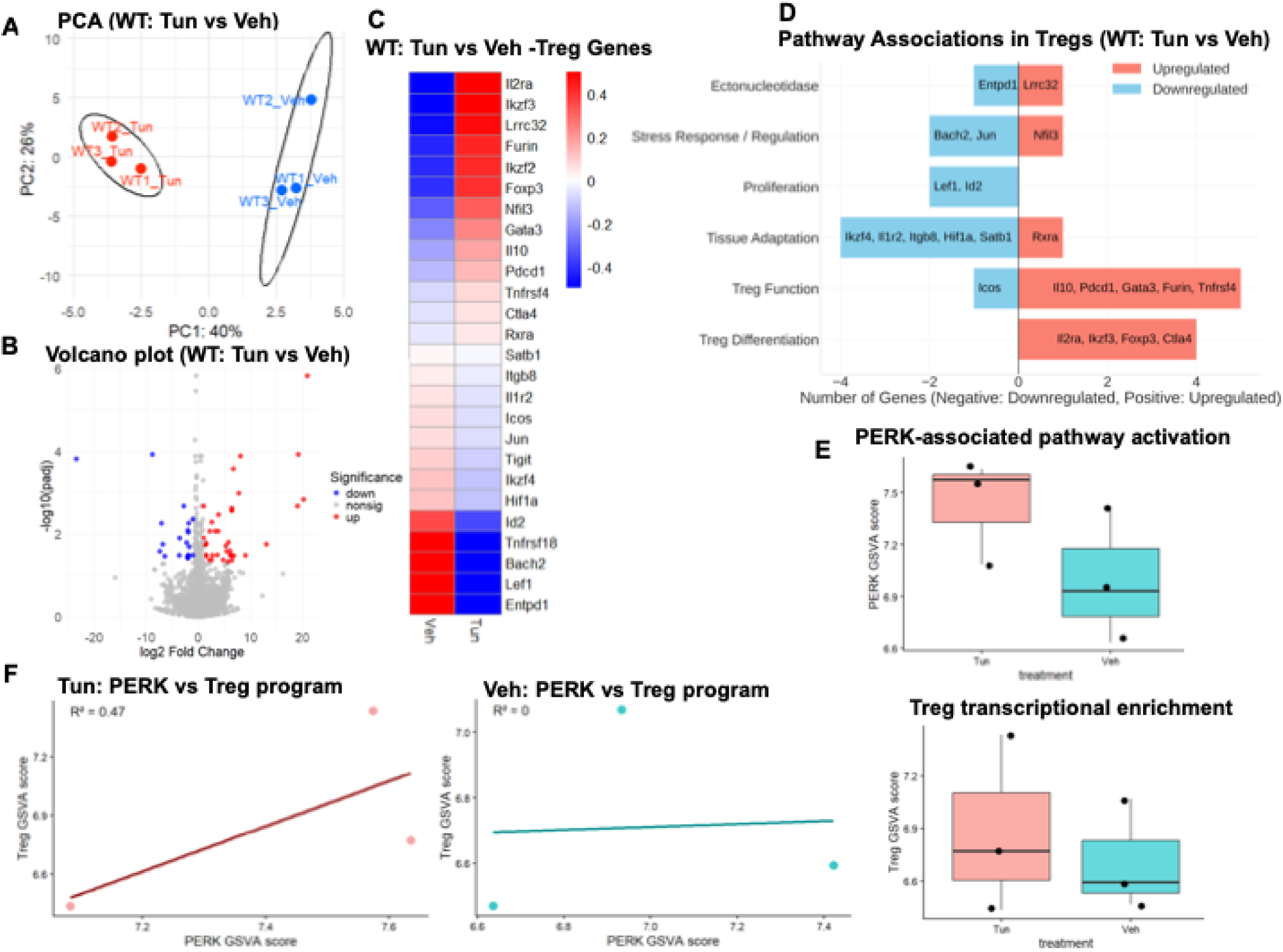
Tunicamycin-induced ER stress is associated with enrichment of Treg transcriptional programs and PERK-associated pathway activity in WT CD4^+^ T cells. **(A)** Principal component analysis (PCA) of bulk RNA-sequencing data from WT CD4^+^ T cells treated with tunicamycin (Tun) or vehicle (Veh). **(B)** Volcano plot showing differential gene expression between Tun- and Veh-treated WT CD4^+^ T cells. Red points indicate significantly upregulated genes, blue points indicate significantly downregulated genes, and gray points indicate genes not meeting the significance threshold. **(C)** Heatmap showing relative expression of selected Treg-associated genes between Veh- and Tun-treated cells. **(D)** Functional categorization of differentially expressed Treg-associated genes into pathways related to Treg differentiation, Treg function, tissue adaptation, proliferation, stress response/regulation, and ectonucleotidase activity. Positive values indicate genes upregulated and negative values indicate genes downregulated in Tun relative to Veh. (E) Gene set variation analysis (GSVA) scores for PERK-associated pathway activity and Treg transcriptional enrichment in Tun- and Veh-treated WT CD4^+^ T cells**. (F)** Relationship between PERK-associated GSVA scores and Treg transcriptional enrichment scores within Tun- and Veh-treated samples. Data represent independent biological replicates.

### 3.3 Genetic deletion of PERK reveals genotype-dependent Treg responses to pharmacologic stress-pathway modulation

To determine whether Treg responses to pharmacologic stress-pathway modulation depend on PERK, we compared CD4^+^ T cells isolated from WT-CD4^cre^ and PERK-CD4^cre^ mice under Veh, Tun, CCT, GSK, and Sal conditions. The expected genotypes were confirmed by PCR (Supplementary Figure 1). Analysis of CD4^+^FoxP3^+^ Tregs revealed a significant genotype x treatment interaction, indicating that the effect of PERK deletion depended on the pharmacologic condition. PERK-CD4^cre^ cells exhibited significantly lower FoxP3^+^ Treg frequencies than WT- CD4^cre^ controls under Veh, Tun, and CCT conditions (Figure 3A). Notably, CCT-treated PERK- deficient cells exhibited a lower FoxP3^+^ Treg frequency than CCT-treated WT cells, consistent with attenuation of the full CCT-associated Treg phenotype in the absence of PERK. In contrast, FoxP3^+^ Treg frequencies were higher in PERK-CD4^cre^ than WT-CD4^cre^ cells under GSK and Sal conditions, demonstrating that the influence of PERK deletion varies according to pharmacologic context. Neuropilin-1 expression within CD4^+^FoxP3^+^ Tregs was consistently reduced in PERK-CD4^cre^ cells relative to WT-CD4^cre^ cells across all treatment conditions (Figure 3B). Tun and CCT increased Neuropilin-1 expression relative to Veh in both genotypes, whereas GSK and Sal reduced Neuropilin-1 expression, with treatment effects differing in magnitude according to genotype. These findings indicate that PERK contributes to FoxP3^+^ Treg differentiation and Neuropilin-1-associated phenotypic features during stress-pathway modulation. Additional phenotypic analysis demonstrated that PERK deletion did not uniformly reduce Treg-associated markers. CTLA-4 and TIGIT expression were consistently increased among PERK-deficient FoxP3^+^ Tregs across treatment conditions, whereas ICOS expression was reduced and showed more context-dependent treatment effects (Supplementary Figure 3A-C). Thus, CD4-specific PERK deletion produces selective remodeling of the Treg phenotype, characterized by reduced FoxP3/Neuropilin-1-associated features together with increased checkpoint-receptor expression.

**FIGURE 3.**
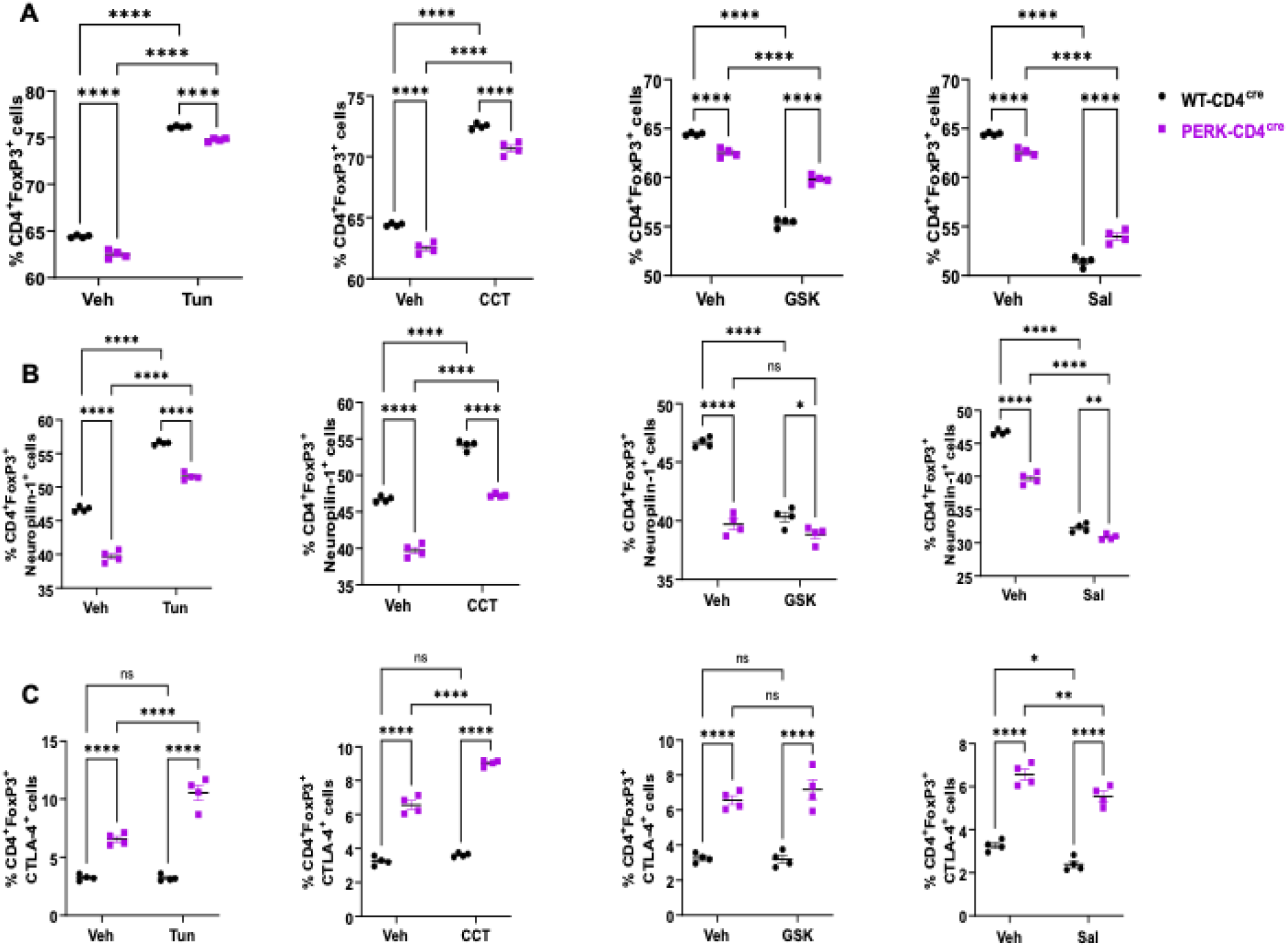
CD4-specific PERK deletion remodels Treg differentiation and regulatory phenotype during pharmacologic stress-pathway modulation. **(A)** Percentage of total CD4^+^ T cells in WT-CD4^cre^ and PERK-CD4^cre^ cultures following treatment with vehicle (Veh), tunicamycin (Tun), CCT020312 (CCT), GSK2606414 (GSK), or salubrinal (Sal). **(B)** Percentage of CD4^+^FoxP3^+^ Tregs across the same treatment conditions. **(C–F)** Percentage of CD4^+^FoxP3^+^ Tregs expressing Neuropilin-1 **(C)**, ICOS **(D)**, CTLA-4 **(E)**, or TIGIT **(F)** in WT-CD4^cre^ and PERK-CD4^cre^ cells. PERK-deficient cells exhibited reduced FoxP3^+^ Treg frequency under Veh, Tun, and CCT conditions and consistently lower Neuropilin-1 expression, whereas CTLA-4 and TIGIT expression were increased across treatment conditions; ICOS showed more limited, treatment-dependent differences. Data are presented as mean ± SEM, with each symbol representing an independent biological replicate. Statistical significance was determined by two-way ANOVA with genotype and treatment as factors followed by Šídák’s multiple-comparisons test. ns, not significant; \**p* < 0.05; \*\**p* < 0.01; \*\*\**p* < 0.001; \*\*\*\**p* < 0.0001.

### 3.4 Pharmacologic stress-pathway modulation alters eIF2α-ATF4 signaling and Treg- associated protein expression

Having established that CD4-specific PERK deletion alters Treg differentiation and phenotype, we next examined how pharmacologic stress-pathway modulation affected PERK/ISR-associated signaling and Treg-associated proteins. To determine how pharmacologic stress-pathway modulation affected PERK/ISR-associated signaling and Treg-associated proteins, we examined p-PERK, p- eIF2α, FoxP3, IL-10, ATF4, and CHOP by immunoblotting (Figure 4A) with densitometric quantifications shown in Figures 4B-G. Across conditions, p-PERK levels showed variability but did not demonstrate statistically significant differences (Figure 4B). In contrast, p-eIF2α was significantly reduced in CCT-treated cells compared with Veh controls at the measured time point, indicating altered downstream stress-signaling (Figure 4C). Notably, the absence of a significant increase in p-PERK together with reduced p-eIF2α at this 4-hour time point does not demonstrate sustained canonical PERK activation by CCT and may reflect temporal regulation or feedback within the integrated stress response. FoxP3 protein abundance was significantly affected by pharmacologic treatment (Figure 4D). Specifically, FoxP3 levels were reduced in GSK-treated cells compared with Tun-treated cells, and Sal treatment also significantly decreased FoxP3 compared with Veh controls. Also, that IL-10 protein levels exhibited modest variation across treatments but did not reach statistical significance (Figure 4E). ATF4 expression was significantly reduced following Sal treatment compared with vehicle (Figure 4F), whereas CHOP levels varied without statistically significant differences (Figure 4G). Collectively, these findings demonstrate that pharmacologic manipulation of PERK/ISR-associated stress signaling is accompanied by changes in downstream stress-response proteins and FoxP3 abundance. The observed effects, particularly those associated with GSK and Sal, should be interpreted in the context of the distinct pharmacologic targets and selectivity of these compounds.

**FIGURE 4.**
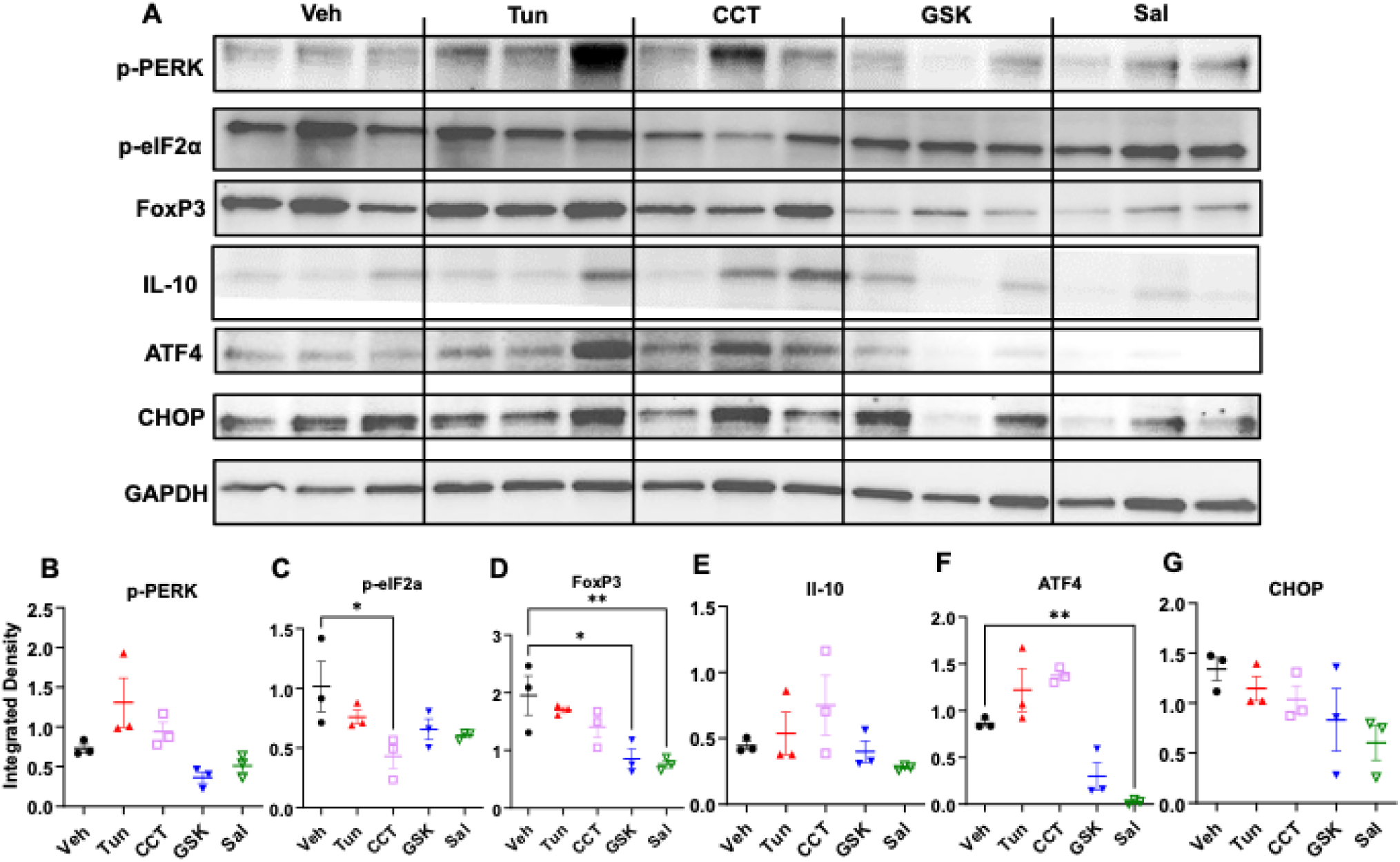
Pharmacologic stress-pathway modulation alters PERK/ISR-associated signaling and Treg-associated protein expression. **(A)** Representative immunoblots showing p-PERK, p- eIF2α, FoxP3, IL-10, ATF4, and CHOP protein levels following treatment with vehicle (Veh), tunicamycin (Tun), CCT020312 (CCT), GSK2606414 (GSK), or salubrinal (Sal). GAPDH served as a loading control. **(B–G)** Densitometric quantification of p-PERK **(B)**, p-eIF2α **(C)**, FoxP3 **(D)**, IL- 10 **(E)**, ATF4 **(F)**, and CHOP **(G)** normalized to GAPDH. Data are presented as mean ± SEM from three independent biological replicates. Statistical analysis was performed using one-way ANOVA followed by Tukey’s multiple-comparisons test. \**p* < 0.05; \*\**p* < 0.01.

### 3.5 Pharmacologic modulation of PERK-related stress signaling alters Treg-mediated control of effector T-cell proliferation

We next evaluated whether pharmacologic modulation of PERK-associated stress signaling influenced the ability of Tregs to control effector T (Teff)-cell proliferation. In the in vitro suppression assay, Teff proliferation was quantified by CellTrace dilution across a range of Treg ratios. CCT treatment was associated with significantly reduced Teff-cell proliferation compared with Veh at several intermediate Treg ratios, consistent with enhanced Treg-mediated suppression (Figure 5A). Representative proliferation histograms at the 1:8 Treg ratio similarly showed lower Teff proliferation in the CCT condition compared with Veh treatment. In contrast, GSK treatment was associated with increased Teff-cell proliferation relative to Veh at several Treg ratios (Figure 5B), consistent with reduced Treg-mediated suppression. Sal treatment similarly increased Teff-cell proliferation, with the most prominent differences observed at intermediate Treg ratios (Figure 5C). Representative histograms at the 1:8 ratio further illustrate the increased proportion of proliferating Teff cells under GSK- and Sal-associated conditions. Together, these findings demonstrate that pharmacologic modulation of PERK/ISR-associated stress signaling alters the capacity of Tregs to restrain Teff-cell proliferation, with CCT-associated conditions favoring greater suppression and GSK-and Sal-associated conditions favoring increased Teff-cell proliferation.

**FIGURE 5.**
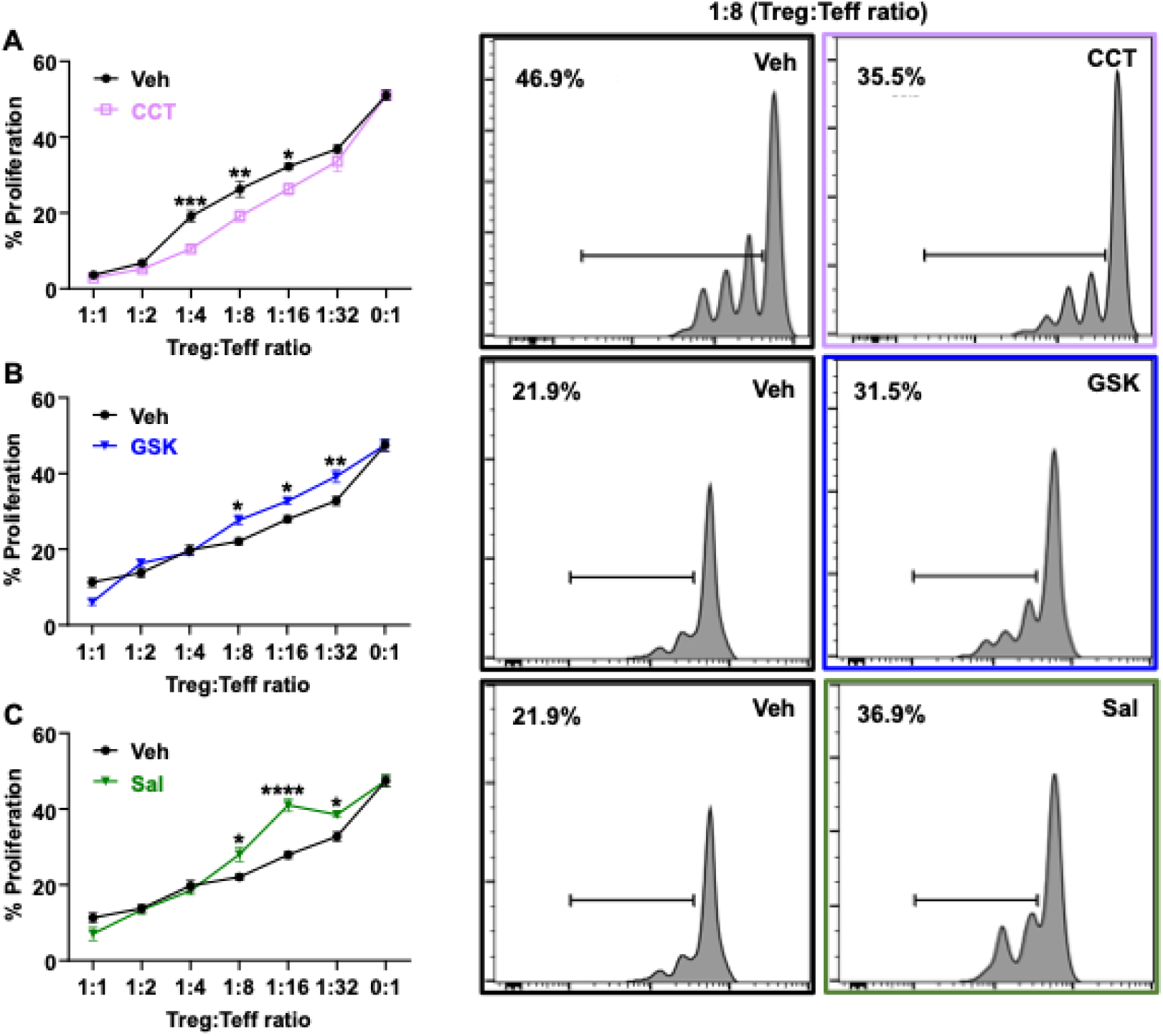
Pharmacologic modulation of PERK/ISR-associated stress signaling alters Treg- mediated control of effector T-cell proliferation. **(A)** Percentage of proliferating effector T (Teff) cells across increasing Treg:Teff ratios under vehicle (Veh)- and CCT020312 (CCT)-associated conditions. CCT treatment was associated with significantly reduced Teff-cell proliferation at several intermediate Treg:Teff ratios, consistent with enhanced Treg-mediated suppression. Representative CellTrace proliferation histograms at the 1:8 Treg:Teff ratio are shown at right. **(B)** Teff-cell proliferation across Treg:Teff ratios under Veh- and GSK2606414 (GSK)-associated conditions. GSK treatment was associated with increased Teff-cell proliferation at several ratios, consistent with reduced Treg-mediated suppression. Representative histograms at the 1:8 ratio are shown at right. **(C)** Teff-cell proliferation under Veh- and salubrinal (Sal)-associated conditions. Sal treatment similarly increased Teff-cell proliferation at several intermediate ratios, consistent with reduced Treg-mediated suppression. Representative histograms at the 1:8 ratio are shown at right. Data are presented as mean ± SEM. Statistical analysis was performed using two-way ANOVA followed by Šídák’s multiple- comparisons test. \**p* < 0.05; \*\**p* < 0.01; \*\*\**p* < 0.001; \*\*\*\**p* < 0.0001.

### 3.6 Single-cell RNA sequencing identifies a PERK-high, stress-adapted Treg state

To examine heterogeneity in PERK-associated stress signaling in vivo, we performed single-cell RNA sequencing of isolated LP cells. UMAP projection revealed distinct segregation of non-Treg and Treg populations based on canonical marker expression (Figure 6A). PERK module scoring demonstrated preferential enrichment of PERK-associated module scores within Tregs compared with non-Tregs (Figure 6B). Violin plot analysis confirmed elevated expression of PERK/eIF2α/ATF4-associated genes, including *Atf4*, *Ddit3*, and *Ppp1r15a*, in Tregs (Figure 6C). Sub-clustering of Tregs identified discrete PERK-low and PERK-high subsets (Figure 6D). PERK- high Tregs exhibited significantly increased module scores and upregulation of canonical stress- adaptive genes relative to PERK-low Tregs. Heatmap analysis further demonstrated enrichment of *Eif2ak3*, *Atf4*, *Ddit3*, *Ppp1r15a*, *Asns*, *Hspa5*, and *Herpud1* in the PERK-high subset (Figure 6E). These findings reveal transcriptional heterogeneity within the Treg compartment and identify a PERK-high stress-adapted Treg state characterized by enrichment of PERK/ISR-associated and broader tissue-adaptive programs.

**FIGURE 6.**
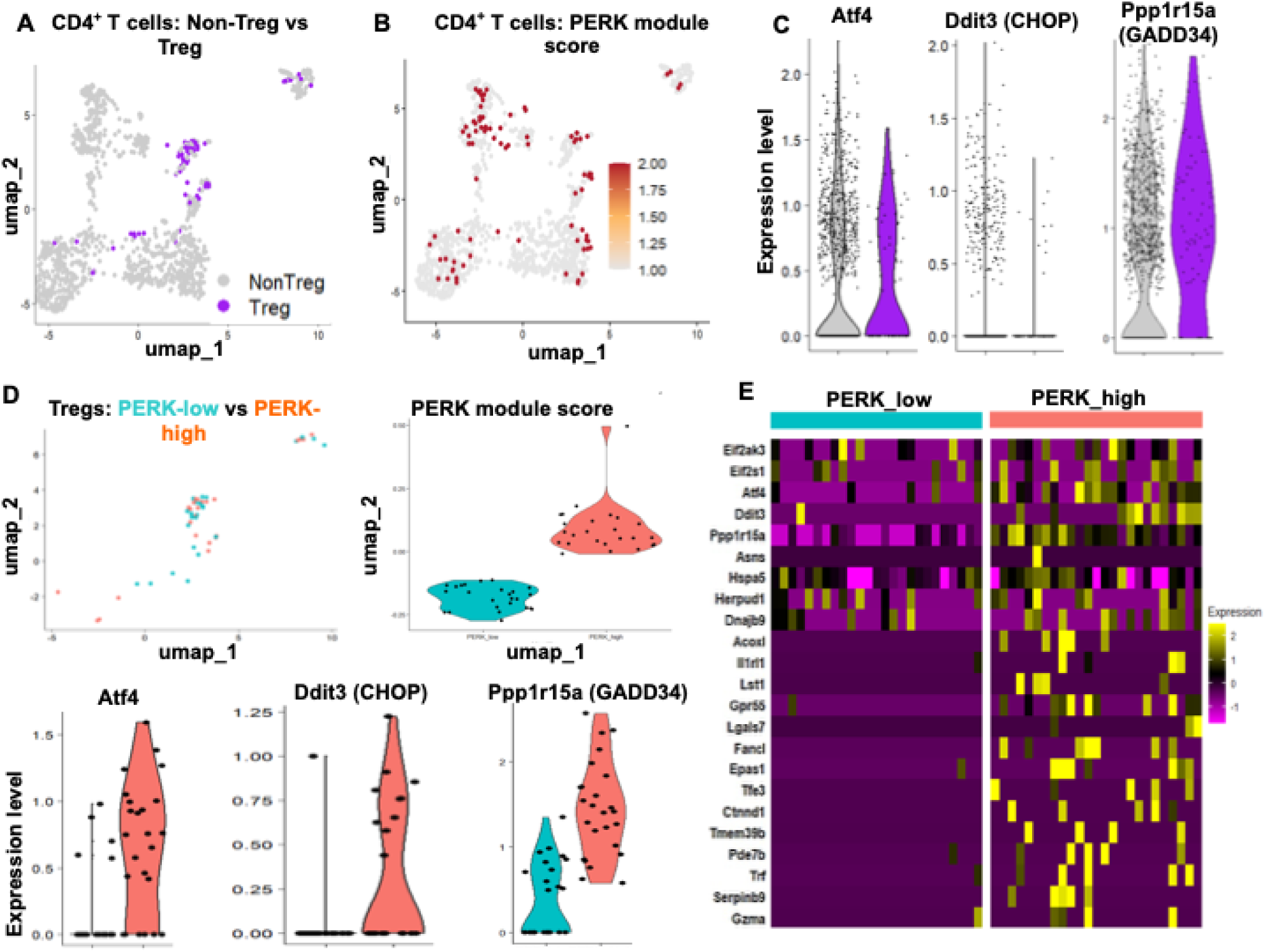
Single-cell RNA sequencing identifies enrichment of PERK-associated stress- response programs and a PERK-high state within intestinal Tregs. **(A)** UMAP projection of CD4^+^ T cells highlighting non-Treg and Treg populations based on canonical marker expression. **(B)** Feature plot showing PERK-associated module scores across CD4^+^ T cells. **(C)** Violin plots comparing expression of ISR-associated genes *Atf4, Ddit3* (CHOP), and *Ppp1r15a* (GADD34) between non-Tregs and Tregs. **(D)** Stratification of Tregs according to PERK-associated module scores identifies PERK-low and PERK-high transcriptional states. UMAP visualization shows the distribution of these populations, while violin plots compare PERK-associated module scores and expression of *Atf4, Ddit3,* and *Ppp1r15a*. **(E)** Heatmap showing differentially expressed genes between PERK-low and PERK-high Tregs, including genes associated with PERK/ISR signaling, proteostasis, metabolic adaptation, and tissue-responsive programs. Single-cell analyses were performed using Seurat in R.

**FIGURE 7.**
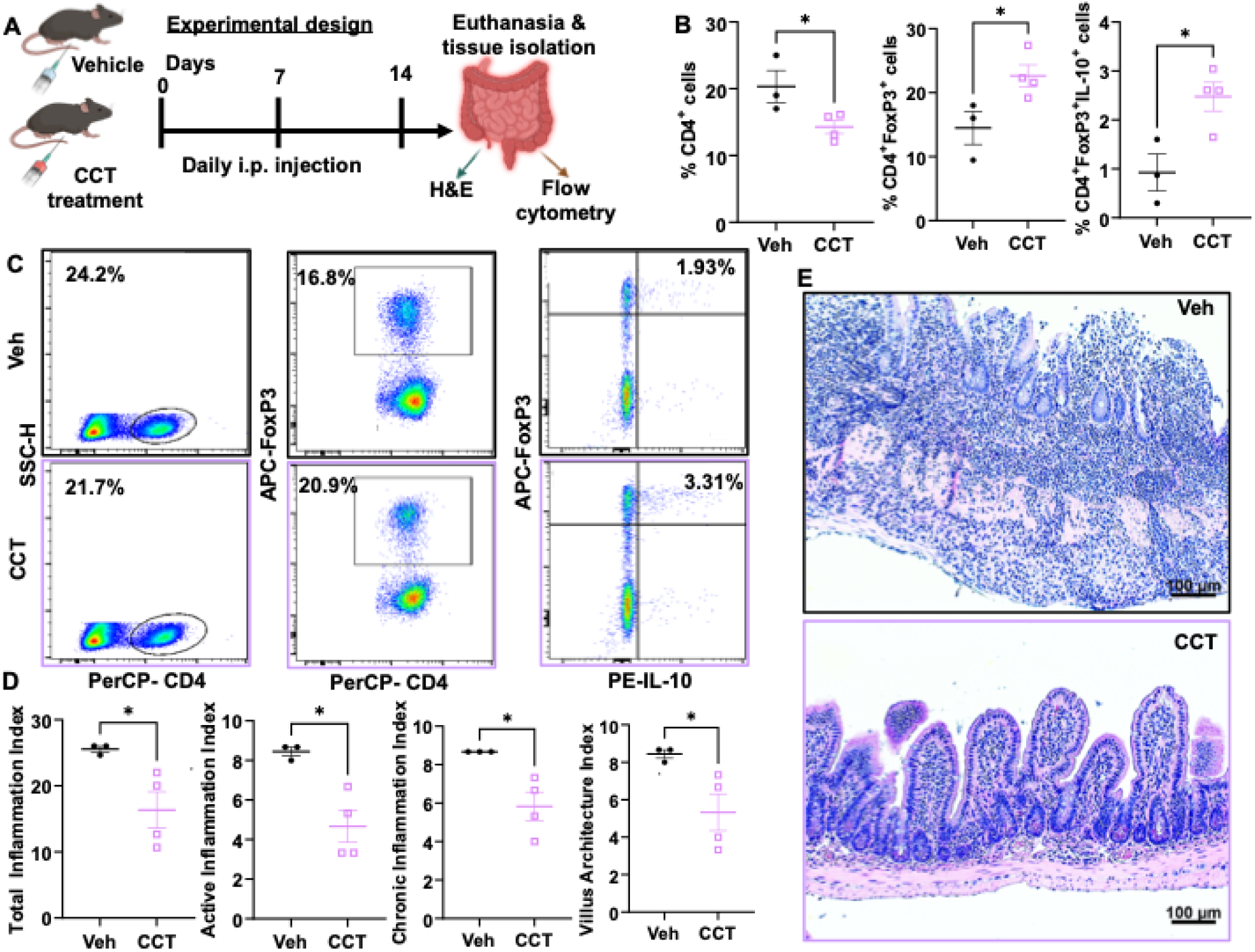
Systemic CCT treatment is associated with increased lamina propria Treg responses and reduced ileal inflammation in TNF^ΔARE^ mice. **(A)** Schematic representation of the in vivo experimental design. Male TNF^ΔARE^ mice (16-18 weeks old) received CCT020312 (CCT; 4 mg/kg) or vehicle (Veh; DMSO diluted in 1× PBS) by daily intraperitoneal injection for 14 consecutive days. On day 14, mice were euthanized and ileal tissue was collected for flow-cytometric and histologic analyses. **(B)** Flow-cytometric quantification of total CD4^+^ cells, CD4^+^FoxP3^+^ Tregs, and IL-10-producing CD4^+^FoxP3^+^ Tregs within ileal lamina propria leukocytes from Veh- and CCT- treated mice. **(C)** Representative flow-cytometry plots showing CD4^+^, CD4^+^FoxP3^+^, and CD4^+^FoxP3^+^IL-10^+^ populations in Veh- and CCT-treated mice. **(D)** Histologic assessment of distal ileal inflammation, including total inflammation, active inflammation, chronic inflammation, and villus architecture indices. **(E)** Representative H&E-stained distal ileal sections from Veh- and CCT- treated TNF^ΔARE^ mice. Scale bars, 100 μm. Data are presented as mean ± SEM; each symbol represents an individual mouse. Statistical significance was determined using an unpaired two-tailed Student’s *t*-test. \**p* < 0.05.

### 3.7 Systemic CCT treatment is associated with increased intestinal Treg responses and reduced ileal inflammation in TNF^ΔARE+/-^ mice

To determine whether systemic CCT treatment alters intestinal Treg responses under inflammatory conditions in vivo, TNF^ΔARE /^ mice were treated with CCT or Veh control for 14 days (Figure 7A). Following treatment, LP leukocytes were isolated from the ileum and analyzed by flow cytometry. Quantitative analysis revealed that CCT- treated mice had a significantly increased proportion of CD4 FoxP3 regulatory T cells within the LP compared with Veh-treated mice (Figure 7B). In addition, the frequency of IL-10-producing CD4 FoxP3 Tregs was significantly elevated in CCT-treated mice. Representative flow cytometry plots illustrate the gating strategy and the corresponding FoxP3 and IL-10 Treg populations in CCT-treated mice (Figure 7C). To assess intestinal inflammation, distal ileal sections were examined histologically using H&E staining. Quantitative histologic scoring demonstrated that CCT-treated TNF^ΔARE /^ mice exhibited significantly reduced ileal inflammation compared with Veh-treated controls across multiple indices, including total inflammation, active inflammation, chronic inflammation, and tissue architecture disruption (Figure 7D). Representative histological images further illustrate improved mucosal architecture and reduced inflammatory infiltrates in CCT-treated mice (Figure 7E). Representative gating strategies for CD8^+^ T cells and FoxP3^neg^ effector CD4^+^ T-cell cytokine populations are shown in Supplementary Figure 2. No significance differences were observed between Veh- and CCT- treated mice in the frequency of CD8^+^ lymphocytes, CD4^+^FoxP3^neg^IFNγ^+^ cells or CD4^+^FoxP3^neg^IL- 17^+^ cells. Collectively, these findings demonstrate that systemic CCT treatment is associated with increased LP Treg responses and reduced ileal inflammation in TNF^ΔARE /^ mice. Since CCT was administered systematically, these experiments do not establish whether the observed changes result from direct effects on Tregs or from modulation of other intestinal immune, epithelial, or stromal cell populations.

## 4 Discussion

While the UPR has been studied extensively as an ER stress adaptation mechanism, the contribution of PERK-associated signaling to Treg biology remains incompletely defined (26, 34–36). Importantly, the present study distinguishes between the UPR and ISR while also highlighting their intersection. The UPR comprises the IRE1, PERK, and ATF6 branches, whereas the ISR is a broader adaptive pathway converging on eIF2α phosphorylation and ATF4 translation in response to multiple stressors. PERK occupies a unique position at this interface by coupling ER stress to ISR activation (9). In this study, complementary pharmacologic and genetic approaches identify PERK as a context-dependent regulator of Treg differentiation and phenotype. Pharmacologic modulation of PERK/ISR-associated signaling altered FoxP3^+^ Treg differentiation, stress-response proteins, and Treg-mediated control of effector T-cell proliferation, while CD4-specific PERK deletion provided genetic evidence that Treg responses to these conditions depend in part on PERK. Notably, PERK deficiency did not uniformly suppress regulatory-associated features but instead produced selective phenotypic remodeling. In vivo, systemic CCT treatment increased LP Treg responses and was associated with reduced ileal inflammation, although these findings do not establish that the anti- inflammatory effect is mediated specifically through CD4^+^ T-cell PERK signaling.

Beyond transcriptional effects, pharmacologic modulation of PERK-associated stress signaling was also accompanied by changes in FoxP3 protein abundance, suggesting a potential additional level of regulation of Treg identity (37, 38). Although IL-10 protein levels did not change significantly by immunoblot, this does not necessarily conflict with the observed changes in FoxP3 abundance and Treg-related function. FoxP3 primarily reflects lineage identity (39, 40), whereas IL- 10 is a context-dependent secreted effector molecule subject to dynamic regulation and assay- dependent variability (41). Thus, pharmacologic modulation of the PERK/ISR stress-response network may influence Treg lineage integrity without uniformly altering all suppressive effector proteins under the same experimental conditions. Moreover, FoxP3 abundance is regulated post- transcriptionally through pathways controlling protein folding, ubiquitin-proteasome turnover, and stress-responsive translation (42, 43). Adaptive PERK-eIF2α-ATF4 signaling has the potential to enhance proteostatic fitness by transiently reducing global translation while enabling selective translation of stress-adaptive programs that support amino acid homeostasis, redox balance, and ER quality control(44, 45). Such changes could favor preservation of FoxP3 protein by reducing proteotoxic burden and limiting degradation pathways engaged during inflammatory or metabolic stress (46, 47). The genetic data extend this pharmacologic observation by demonstrating reduced FoxP3^+^ Treg frequency in PERK-deficient cells under basal, Tun, and CCT conditions. Although these findings support a contribution of PERK to the maintenance or induction of the FoxP3^+^ Treg phenotype, direct regulation of FoxP3 protein stability by PERK will require dedicated turnover or lineage-stability studies.

The differential effects of CD4-specific PERK deletion across Treg-associated markers further indicate that PERK does not uniformly regulate all components of the regulatory program. The principal changes highlighted in the main analysis were reduced FoxP3^+^ Treg frequency and reduced Neuropilin-1 expression following PERK deletion. Complementary phenotyping demonstrated increased CTLA-4 and TIGIT expression and context-dependent changes in ICOS among PERK- deficient FoxP3^+^ Tregs (Supplementary Figure 3). FoxP3 and Neuropilin-1 are associated with Treg lineage identity, stability, and tissue adaptation (48–51), whereas CTLA-4 and TIGIT represent distinct inhibitory mechanisms that contribute to suppressive activity (52–54), and ICOS supports Treg activation, survival, and functional adaptation (17, 55). Accordingly, these findings may reflect remodeling of the Treg compartment rather than generalized loss of regulatory function. One possibility is that impaired PERK-dependent stress adaptation compromises maintenance of canonical FoxP3/Neuropilin-1-associated features while potentially promoting compensatory upregulation of inhibitory receptors. Alternatively, PERK deficiency may favor the persistence or expansion of a checkpoint-high Treg state. Because individual Tregs can co-express several of these molecules (56, 57), the current single-marker analyses cannot determine whether these changes occur within the same cells or reflect shifts among phenotypically distinct Treg states. Therefore, functional and multidimensional co-expression studies will be required to determine whether increased CTLA-4 and TIGIT compensate for, or merely accompany, reduced canonical Treg-associated features.

These findings align with emerging evidence that adaptive ER stress signaling and ISR engagement support immune tolerance by maintaining proteostatic and metabolic fitness (11, 15, 23). Tregs operate in metabolically demanding and inflammatory microenvironments, particularly in barrier tissues like the intestine (23). Prior studies have shown that UPR activity can support Treg survival and IL-10 production under stress conditions (26, 58). The current data extend these observations by showing that pharmacologic modulation of PERK-associated stress signaling alters Treg frequency, expression of immune regulatory receptors, including CTLA-4, TIGIT, ICOS, Neuropilin-1, and the capacity of Tregs to restrain effector T-cell proliferation. Transcriptomic profiling further revealed coordinated enrichment of core Treg-related genes, including *Foxp3*, *Il2ra*, *Il10*, and *Ikzf2*, together with PERK-associated pathway signatures. These transcriptional findings were accompanied by treatment-associated changes in FoxP3 protein abundance. Collectively, the pharmacologic, transcriptional, and genetic observations support a model in which PERK/ISR- associated stress signaling contributes to Treg adaptation by linking cellular stress responses with regulatory transcriptional and translational programs. The divergent marker responses observed following PERK deletion further suggest that this regulation is multidimensional rather than uniformly directed toward either gain or loss of Treg identity.

However, excessive or chronic ER stress can impair Treg stability or favor inflammatory polarization, particularly toward Th17 differentiation (22, 59). Sustained UPR activation has been linked to CHOP-mediated apoptosis, oxidative stress, and FoxP3 destabilization in inflammatory contexts (23, 60). Additionally, IRE1α RNase activity can degrade transcripts and microRNAs that influence Treg maintenance (6, 22, 61). These findings underscore that UPR signaling outcomes depend on the intensity, duration, and branch-specific nature of the stress response. In the present study, the pharmacologic conditions associated with enhanced Treg responses were not accompanied by a significant increase in CHOP expression, suggesting that these treatments did not induce a dominant CHOP-associated maladaptive stress response at the measured time point. Together, the data are consistent with the possibility that controlled modulation of PERK/ISR-associated signaling may favor an adaptive stress-response state that supports Treg-associated phenotypes. The genetic data further demonstrate that PERK contributes to these responses, although the persistence of treatment-associated effects in PERK-deficient cells indicates that additional UPR and ISR pathways are also likely involved.

Single-cell RNA sequencing identified a distinct PERK-high Treg subset enriched for canonical PERK-associated genes, including *Eif2ak3*, *Atf4*, *Ddit3*, *Ppp1r15a*, *Asns*, *Hspa5*, and *Herpud1*. Beyond these core stress-response genes, the heatmap (Figure 6E) also revealed enrichment of transcripts linked to broader environmental adaptation, including hypoxic and metabolic regulation (*Epas1*), tissue-responsive or activated immune states (*Il1rl1*, *Gpr55*), stress-responsive transcriptional control (*Tfe3*), and cytoprotective or effector-associated programs (*Serpinb9*, *Gzma*). Together, these findings suggest that PERK-high Tregs do not simply reflect ER stress engagement alone, but rather a broader stress-adapted regulatory state shaped by inflammatory, metabolic, and tissue-derived cues. Thus, enrichment of PERK-associated transcriptional programs appears to mark functional heterogeneity within the Treg compartment. Rather than indicating a dysfunctional state, PERK-high Tregs exhibited a transcriptional profile consistent with adaptive proteostasis and metabolic remodeling (26). These observations suggest that PERK-associated transcriptional activity marks a stress-adapted intestinal Treg state and, in conjunction with the genetic findings, is consistent with a contribution of PERK signaling to Treg heterogeneity. However, whether the PERK-high transcriptional state is functionally related to the phenotypic changes observed following PERK deletion remains to be determined.

The in vivo TNF^ΔARE+/-^ model further supports an association between pharmacologic modulation of PERK-associated stress signaling and regulatory immune responses during intestinal inflammation. Systemic CCT treatment increased LP Treg frequency and the proportion of IL-10- producing Tregs while reducing histologic inflammation, consistent with enhanced regulatory responses in chronically inflamed intestinal tissue. TNF^ΔARE+/-^ mice experience persistent inflammatory signaling and metabolic stress (62), providing a relevant setting in which adaptive stress-response pathways may influence immune-cell function. However, because CCT was administered systematically, these findings do not establish that the reduction in inflammation was mediated directly through Treg-intrinsic PERK signaling. Effects on epithelial, myeloid, stromal, or other immune-cell populations could also contribute to the observed changes in Treg frequency and intestinal pathology. Therefore, the in vivo data support a relationship between CCT treatment, increased regulatory immune responses, and reduced ileal inflammation, while the cell-intrinsic contribution of PERK signaling to these effects remains to be defined. One caveat is that the ex vivo genetic findings strengthen the plausibility that CD4^+^ T-cell PERK signaling contributes to regulatory responses; however, they do not establish that the anti-inflammatory effects of systemic CCT treatment in TNF^ΔARE+/-^ mice are mediated specifically through this compartment.

Transcriptomic analysis suggested a potential trade-off between regulatory and proliferative programs, with enrichment of Treg-associated suppressive and lineage-defining genes occurring alongside reduced expression of pathways related to proliferation and tissue adaptation. This pattern is consistent with the possibility that PERK/ISR-associated stress signaling may favor a more differentiated regulatory state while limiting transcriptional programs associated with expansion or environmental plasticity. In settings requiring rapid Treg proliferation, sustained engagement of stress-response pathways could therefore constrain proliferative flexibility despite supporting regulatory-associated features. Conversely, pharmacologic conditions associated with reduced PERK/ISR signaling were accompanied by impaired control of effector T-cell proliferation, although these findings do not establish a direct causal relationship between PERK activity and Treg proliferative capacity.

Pharmacologic specificity remains an important consideration because the compounds used in this study differ in potency, selectivity, and mechanism of action. Sal acts indirectly by modulating eIF2α dephosphorylation rather than directly inhibiting PERK (63–65), while Tun and some pharmacologic PERK activators can engage broader ER stress pathways beyond PERK alone (66–68). The present genetic experiments address this limitation in part: CD4-specific PERK deletion significantly altered Treg responses across pharmacologic conditions and attenuated the FoxP3^+^ Treg phenotype associated with CCT treatment. Nevertheless, CCT-associated responses were not completely abolished in PERK-deficient cells, and the effects of GSK and Sal differed substantially according to genotype. These findings indicate that PERK contributes to the observed pharmacologic response while also supporting involvement of PERK-independent ISR signaling and crosstalk with the IRE1α and ATF6 branches of the UPR. Thus, pharmacologic and genetic perturbation should be considered complementary rather than interchangeable approaches to defining PERK-dependent Treg biology.

## 5 Conclusion

Collectively, our findings support a model in which PERK-associated/ISR stress signaling contributes to Treg differentiation, phenotype, and functional adaptation under ER-stress and inflammatory conditions. Importantly, CD4-specific PERK deletion provides genetic evidence that PERK contributes to basal and pharmacologically induced Treg responses, including the full FoxP3^+^ Treg phenotype associated with CCT treatment. The divergent effects of PERK deletion on FoxP3 and Neuropilin-1 versus CTLA-4 and TIGIT further demonstrate that PERK does not uniformly regulate the Treg program but instead shapes distinct regulatory features in a context-dependent manner. Together with the identification of a PERK-high stress-adapted intestinal Treg state and the association of systemic CCT treatment with increased intestinal Treg responses and reduced ileitis, these findings identify PERK as an important component of the broader stress-response network governing regulatory immunity. Further studies using Treg-specific genetic models and direct functional assays will be required to define PERK-dependent mechanisms within mature Tregs and determine their therapeutic relevance in intestinal inflammation.

## 6 Limitations

Several limitations remain. First, CD4-Cre-mediated Eif2ak3 deletion provides evidence for a CD4^+^ T-cell-intrinsic contribution of PERK but is not specific to mature FoxP3^+^ Tregs; developmental or conventional CD4^+^ T-cell effects therefore cannot be excluded. Second, the genetic experiments define phenotypic changes but do not directly establish the suppressive capacity of PERK-deficient Tregs. Future studies using Foxp3-directed Eif2ak3 deletion and genotype-specific suppression assays will be important for distinguishing Treg-intrinsic mechanisms from effects occurring earlier in CD4^+^ T-cell differentiation. Third, systemic CCT treatment may influence multiple intestinal cell populations, and therefore the in vivo anti-inflammatory response cannot be attributed exclusively to Tregs.

## 7 Data availability statement

The bulk RNA-seq and single-cell RNA-seq data generated in this study are available in the GEO repository under accession number GSE313127, and GSE320371, respectively. The R code used for data processing and analysis is available upon request and can be shared via GitHub.

## 9 Author Contributions

KMA: Conceptualization, Data curation, Formal analysis, Investigation, Methodology, Project administration, Software, Validation, Visualization, Writing - original draft, Writing - review & editing. TTN: Investigation, Methodology, Validation, Writing - review & editing. NB: Investigation, Writing - review & editing. TP: Resources, Software, Writing - review & editing. GRK: Investigation, Writing - review & editing. PJ: Investigation, Writing - review & editing. EFZ: Conceptualization, Formal analysis, Funding acquisition, Resources, Project administration, Methodology, Supervision, Validation, Visualization, Writing - review & editing.

## 11 Generative AI statement

The authors declare that Google Gemini was used for editing some sentence structures, and reference search.

---

**Supplementary Figure 1.**
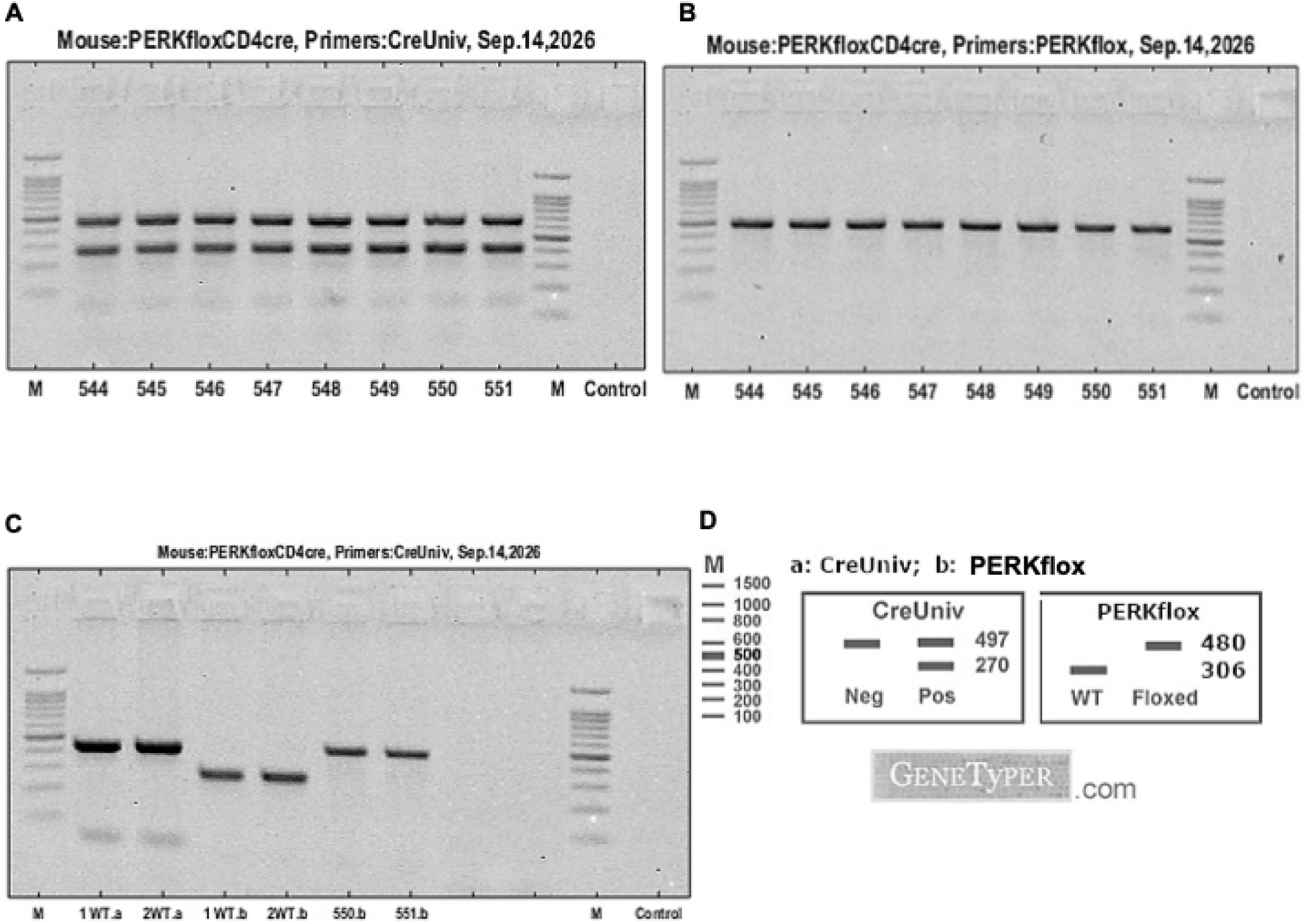
Genotyping validation of CD4-specific PERK conditional knockout mice. **(A)** PCR genotyping using Cre universal primers showing the expected Cre-positive banding pattern in mice 544–551. **(B)** PCR genotyping using PERKflox primers confirming the presence of the floxed PERK allele in the same mice. **(C)** Representative genotyping of selected WT-CD4^cre^ and PERK-CD4^cre^ samples using Cre universal and PERKflox primer sets, demonstrating the expected genotype-specific banding patterns. **(D)** Reference schematic indicating the expected PCR product sizes for the Cre universal and PERKflox assays. Expected Cre universal products are 497 bp and 270 bp for Cre-positive samples, whereas the PERKflox assay distinguishes WT and floxed alleles at approximately 306 bp and 480 bp, respectively.

**Supplementary Figure 2.:**
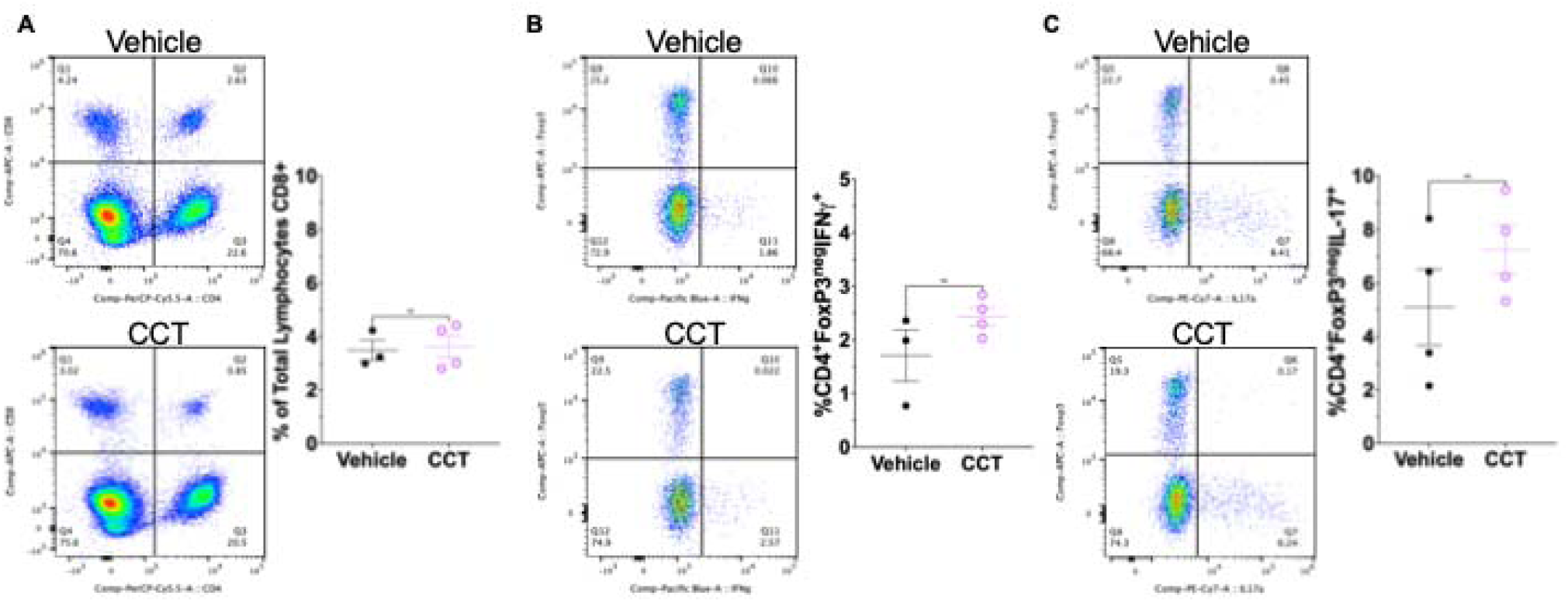

## References

1. Kettel P, Karagoz GE. Endoplasmic Reticulum: Monitoring and Maintaining Protein and Membrane Homeostasis in the Endoplasmic Reticulum by the Unfolded Protein Response. Int J Biochem Cell Biol (2024) 172:106598. Epub 20240518. doi: 10.1016/j.biocel.2024.106598.

2. Coleman OI, Haller D. Er Stress and the Upr in Shaping Intestinal Tissue Homeostasis and Immunity. Front Immunol (2019) 10:2825. Epub 20191204. doi: 10.3389/fimmu.2019.02825.

3. Read A, Schroder M. The Unfolded Protein Response: An Overview. Biology (Basel) (2021) 10(5). Epub 20210429. doi: 10.3390/biology10050384.

4. Wong MY, DiChiara AS, Suen PH, Chen K, Doan ND, Shoulders MD. Adapting Secretory Proteostasis and Function through the Unfolded Protein Response. Curr Top Microbiol Immunol (2018) 414:1–25. doi: 10.1007/82_2017_56.

5. Gawlak-Socka S, Kowalczyk E, Wiktorowska-Owczarek A. Unfolded Protein Response at the Crossroads: Integrating Endoplasmic Reticulum Stress with Cellular Stress Networks. Int J Mol Sci (2026) 27(4). Epub 20260219. doi: 10.3390/ijms27041986.

6. Duan J, Huang D, Fong YW. Targeting the Upr with Small Molecules: Emerging Strategies for Immune Regulation. Molecules (2026) 31(3). Epub 20260205. doi: 10.3390/molecules31030559.

7. Costa-Mattioli M, Walter P. The Integrated Stress Response: From Mechanism to Disease. Science (2020) 368(6489). doi: 10.1126/science.aat5314.

8. Pakos-Zebrucka K, Koryga I, Mnich K, Ljujic M, Samali A, Gorman AM. The Integrated Stress Response. EMBO Rep (2016) 17(10):1374–95. Epub 20160914. doi: 10.15252/embr.201642195.

9. Perea V, Baron KR, Dolina V, Aviles G, Kim G, Rosarda JD, et al. Pharmacologic Activation of a Compensatory Integrated Stress Response Kinase Promotes Mitochondrial Remodeling in Perk- Deficient Cells. Cell Chem Biol (2023) 30(12):1571–84 e5. Epub 20231102. doi: 10.1016/j.chembiol.2023.10.006.

10. Moreews M, Karlsson MCI. Endoplasmic Reticulum Stress: A Key Player in Immune Cell Regulation and Autoimmune Disorders. Semin Immunol (2025) 78:101954. Epub 20250422. doi: 10.1016/j.smim.2025.101954.

11. Fu Z, Li M, Zhou H, Zhong X, Zhang Z, Meng X, et al. Endoplasmic Reticulum Stress Orchestrates Tumor Metabolism and Immunity: New Insights into Immunometabolic Therapeutics. Front Immunol (2025) 16:1674163. Epub 20250930. doi: 10.3389/fimmu.2025.1674163.

12. Nghiem TT, Nguyen KA, Kusuma F, Park S, Park J, Joe Y, et al. The Perk-Eif2alpha-Atf4 Axis Is Involved in Mediating Er-Stress-Induced Ferroptosis Via Ddit4-Mtorc1 Inhibition and Acetaminophen-Induced Hepatotoxicity. Antioxidants (Basel) (2025) 14(3). Epub 20250303. doi: 10.3390/antiox14030307.

13. Raines LN, Zhao H, Wang Y, Chen HY, Gallart-Ayala H, Hsueh PC, et al. Perk Is a Critical Metabolic Hub for Immunosuppressive Function in Macrophages. Nat Immunol (2022) 23(3):431–45. Epub 20220228. doi: 10.1038/s41590-022-01145-x.

14. Mendes A, Gigan JP, Rodriguez Rodrigues C, Choteau SA, Sanseau D, Barros D, et al. Proteostasis in Dendritic Cells Is Controlled by the Perk Signaling Axis Independently of Atf4. Life Sci Alliance (2021) 4(2). Epub 20201221. doi: 10.26508/lsa.202000865.

15. Zhang W, Cao X. Unfolded Protein Responses in T Cell Immunity. Front Immunol (2024) 15:1515715. Epub 20250108. doi: 10.3389/fimmu.2024.1515715.

16. Fiyouzi T, Pelaez-Prestel HF, Reyes-Manzanas R, Lafuente EM, Reche PA. Enhancing Regulatory T Cells to Treat Inflammatory and Autoimmune Diseases. Int J Mol Sci (2023) 24(9). Epub 20230425. doi: 10.3390/ijms24097797.

17. Wang L, Liang Y, Zhao C, Ma P, Zeng S, Ju D, et al. Regulatory T Cells in Homeostasis and Disease: Molecular Mechanisms and Therapeutic Potential. Signal Transduct Target Ther (2025) 10(1):345. Epub 20251014. doi: 10.1038/s41392-025-02326-4.

18. Paterson AM, Lovitch SB, Sage PT, Juneja VR, Lee Y, Trombley JD, et al. Deletion of Ctla-4 on Regulatory T Cells During Adulthood Leads to Resistance to Autoimmunity. J Exp Med (2015) 212(10):1603–21. Epub 20150914. doi: 10.1084/jem.20141030.

19. Chaudhry A, Samstein RM, Treuting P, Liang Y, Pils MC, Heinrich JM, et al. Interleukin-10 Signaling in Regulatory T Cells Is Required for Suppression of Th17 Cell-Mediated Inflammation. Immunity (2011) 34(4):566–78. doi: 10.1016/j.immuni.2011.03.018.

20. Golzari-Sorkheh M, Zuniga-Pflucker JC. Development and Function of Foxp3+ Regulators of Immune Responses. Clin Exp Immunol (2023) 213(1):13–22. doi: 10.1093/cei/uxad048.

21. Morawski PA, Mehra P, Chen C, Bhatti T, Wells AD. Foxp3 Protein Stability Is Regulated by Cyclin-Dependent Kinase 2. J Biol Chem (2013) 288(34):24494–502. Epub 20130712. doi: 10.1074/jbc.M113.467704.

22. Xu Y, Melo-Cardenas J, Zhang Y, Gau I, Wei J, Montauti E, et al. The E3 Ligase Hrd1 Stabilizes Tregs by Antagonizing Inflammatory Cytokine-Induced Er Stress Response. JCI Insight (2019) 4(5). Epub 20190307. doi: 10.1172/jci.insight.121887.

23. Alula KM, Nguyen TT, de Zoeten EF. Regulation of Intestinal Regulatory T Cells Via Stress Response Pathways in Inflammatory Bowel Disease. Autoimmunity (2026) 59(1):2602715. Epub 20251222. doi: 10.1080/08916934.2025.2602715.

24. Cao SS, Zimmermann EM, Chuang BM, Song B, Nwokoye A, Wilkinson JE, et al. The Unfolded Protein Response and Chemical Chaperones Reduce Protein Misfolding and Colitis in Mice. Gastroenterology (2013) 144(5):989–1000 e6. Epub 20130118. doi: 10.1053/j.gastro.2013.01.023.

25. Zhang HS, Chen Y, Fan L, Xi QL, Wu GH, Li XX, et al. The Endoplasmic Reticulum Stress Sensor Ire1alpha in Intestinal Epithelial Cells Is Essential for Protecting against Colitis. J Biol Chem (2015) 290(24):15327–36. Epub 20150429. doi: 10.1074/jbc.M114.633560.

26. Feng ZZ, Luo N, Liu Y, Hu JN, Ma T, Yao YM. Er Stress and Its Perk Branch Enhance Tcr- Induced Activation in Regulatory T Cells. Biochem Biophys Res Commun (2021) 563:8–14. Epub 20210528. doi: 10.1016/j.bbrc.2021.05.061.

27. Wu D, Liang J. Activating Transcription Factor 4: A Regulator of Stress Response in Human Cancers. Front Cell Dev Biol (2024) 12:1370012. Epub 20240327. doi: 10.3389/fcell.2024.1370012.

28. Franco A, Almanza G, Burns JC, Wheeler M, Zanetti M. Endoplasmic Reticulum Stress Drives a Regulatory Phenotype in Human T-Cell Clones. Cell Immunol (2010) 266(1):1–6. Epub 20100924. doi: 10.1016/j.cellimm.2010.09.006.

29. Guan BJ, van Hoef V, Jobava R, Elroy-Stein O, Valasek LS, Cargnello M, et al. A Unique Isr Program Determines Cellular Responses to Chronic Stress. Mol Cell (2017) 68(5):885–900 e6. doi: 10.1016/j.molcel.2017.11.007.

30. Laukova M, Glatman Zaretsky A. Regulatory T Cells as a Therapeutic Approach for Inflammatory Bowel Disease. Eur J Immunol (2023) 53(2):e2250007. Epub 20230106. doi: 10.1002/eji.202250007.

31. Hu Y, Yang Y, Li Y, Zhang Q, Zhang W, Jia J, et al. Th17/Treg Imbalance in Inflammatory Bowel Disease: Immunological Mechanisms and Microbiota-Driven Regulation. Front Immunol (2025) 16:1651063. Epub 20251008. doi: 10.3389/fimmu.2025.1651063.

32. Collins CB, Nguyen TT, Leddy RS, Alula KM, Yeckes AR, Strassheim D, et al. Heat Shock Factor 1 Drives Regulatory T-Cell Induction to Limit Murine Intestinal Inflammation. Mucosal Immunol (2024) 17(1):94–110. Epub 20231107. doi: 10.1016/j.mucimm.2023.11.003.

33. Vidrich A, Buzan JM, Barnes S, Reuter BK, Skaar K, Ilo C, et al. Altered Epithelial Cell Lineage Allocation and Global Expansion of the Crypt Epithelial Stem Cell Population Are Associated with Ileitis in Samp1/Yitfc Mice. Am J Pathol (2005) 166(4):1055–67. doi: 10.1016/S0002-9440(10)62326-7.

34. Cheng Q, Choi HJ, Wu Y, Yuan X, Pugel A, Tian L, et al. Er Stress Sensor Perk Promotes T Cell Pathogenicity in Gvhd by Regulating Er-Associated Degradation. J Clin Invest (2025) 135(23). Epub 20250930. doi: 10.1172/JCI190958.

35. Li A, Song NJ, Riesenberg BP, Li Z. The Emerging Roles of Endoplasmic Reticulum Stress in Balancing Immunity and Tolerance in Health and Diseases: Mechanisms and Opportunities. Front Immunol (2019) 10:3154. Epub 20200211. doi: 10.3389/fimmu.2019.03154.

36. Marciniak SJ, Chambers JE, Ron D. Pharmacological Targeting of Endoplasmic Reticulum Stress in Disease. Nat Rev Drug Discov (2022) 21(2):115–40. Epub 20211026. doi: 10.1038/s41573-021-00320-3.

37. Chang H, Wang Y, Gao L, Zheng SJ. Beyond Conventional Transcriptional Regulation Function: Foxp3 as an Integrative Hub for Chromatin Interactions and Protein Complexes in Immune Regulation. Biology (Basel) (2026) 15(3). Epub 20260130. doi: 10.3390/biology15030254.

38. Luckerbauer B, Villarino AV, Laurence A, Hirahara K, Sciume G, Mikami Y, et al. Preserved Suppressive Function Despite Loss of Foxp3: Insights into the Identity of Regulatory T Cells. bioRxiv (2025). Epub 20250122. doi: 10.1101/2025.01.17.633544.

39. Attias M, Al-Aubodah T, Piccirillo CA. Mechanisms of Human Foxp3(+) T(Reg) Cell Development and Function in Health and Disease. Clin Exp Immunol (2019) 197(1):36–51. Epub 20190401. doi: 10.1111/cei.13290.

40. Xie X, Stubbington MJ, Nissen JK, Andersen KG, Hebenstreit D, Teichmann SA, et al. The Regulatory T Cell Lineage Factor Foxp3 Regulates Gene Expression through Several Distinct Mechanisms Mostly Independent of Direct DNA Binding. PLoS Genet (2015) 11(6):e1005251. Epub 20150624. doi: 10.1371/journal.pgen.1005251.

41. Grover P, Goel PN, Greene MI. Regulatory T Cells: Regulation of Identity and Function. Front Immunol (2021) 12:750542. Epub 20211005. doi: 10.3389/fimmu.2021.750542.

42. van Loosdregt J, Fleskens V, Fu J, Brenkman AB, Bekker CP, Pals CE, et al. Stabilization of the Transcription Factor Foxp3 by the Deubiquitinase Usp7 Increases Treg-Cell-Suppressive Capacity. Immunity (2013) 39(2):259–71. doi: 10.1016/j.immuni.2013.05.018.

43. Deng G, Song X, Fujimoto S, Piccirillo CA, Nagai Y, Greene MI. Foxp3 Post-Translational Modifications and Treg Suppressive Activity. Front Immunol (2019) 10:2486. Epub 20191018. doi: 10.3389/fimmu.2019.02486.

44. Rozpedek W, Pytel D, Mucha B, Leszczynska H, Diehl JA, Majsterek I. The Role of the Perk/Eif2alpha/Atf4/Chop Signaling Pathway in Tumor Progression During Endoplasmic Reticulum Stress. Curr Mol Med (2016) 16(6):533–44. doi: 10.2174/1566524016666160523143937.

45. Wek RC. Role of Eif2alpha Kinases in Translational Control and Adaptation to Cellular Stress. Cold Spring Harb Perspect Biol (2018) 10(7). Epub 20180702. doi: 10.1101/cshperspect.a032870.

46. Lu L, Barbi J, Pan F. The Regulation of Immune Tolerance by Foxp3. Nat Rev Immunol (2017) 17(11):703–17. Epub 20170731. doi: 10.1038/nri.2017.75.

47. Luo Z, Zhang Y, Saleh QW, Zhang J, Zhu Z, Tepel M. Metabolic Regulation of Forkhead Box P3 Alternative Splicing Isoforms and Their Impact on Health and Disease. Front Immunol (2023) 14:1278560. Epub 20231006. doi: 10.3389/fimmu.2023.1278560.

48. Junius S, Mavrogiannis AV, Lemaitre P, Gerbaux M, Staels F, Malviya V, et al. Unstable Regulatory T Cells, Enriched for Naive and Nrp1(Neg) Cells, Are Purged after Fate Challenge. Sci Immunol (2021) 6(61). doi: 10.1126/sciimmunol.abe4723.

49. Yadav M, Louvet C, Davini D, Gardner JM, Martinez-Llordella M, Bailey-Bucktrout S, et al. Neuropilin-1 Distinguishes Natural and Inducible Regulatory T Cells among Regulatory T Cell Subsets in Vivo. J Exp Med (2012) 209(10):1713–22, S1-19. Epub 20120910. doi: 10.1084/jem.20120822.

50. Hori S. Stability of Regulatory T-Cell Lineage. Adv Immunol (2011) 112:1–24. doi: 10.1016/B978-0-12-387827-4.00001-2.

51. Delgoffe GM, Woo SR, Turnis ME, Gravano DM, Guy C, Overacre AE, et al. Stability and Function of Regulatory T Cells Is Maintained by a Neuropilin-1-Semaphorin-4a Axis. Nature (2013) 501(7466):252-6. Epub 20130804. doi: 10.1038/nature12428.

52. Adam KR, Suhail A, Kuchroo VK, Vignali DAA. Inhibitory Receptor Agonists: Emerging Strategies in Immune Modulation. J Exp Med (2026) 223(4). Epub 20260331. doi: 10.1084/jem.20241217.

53. Cui H, Hamad M, Elkord E. Tigit in Cancer: From Mechanism of Action to Promising Immunotherapeutic Strategies. Cell Death Dis (2025) 16(1):664. Epub 20250901. doi: 10.1038/s41419-025-07984-4.

54. Rudd CE. Ctla-4 Co-Receptor Impacts on the Function of Treg and Cd8+ T-Cell Subsets. Eur J Immunol (2009) 39(3):687–90. doi: 10.1002/eji.200939261.

55. Li DY, Xiong XZ. Icos(+) Tregs: A Functional Subset of Tregs in Immune Diseases. Front Immunol (2020) 11:2104. Epub 20200828. doi: 10.3389/fimmu.2020.02104.

56. Sato Y, Osada E, Manome Y. Full-Length Transcriptome Sequencing Reveals Treg-Specific Isoform Expression Upon Activation. Int J Mol Sci (2025) 26(13). Epub 20250630. doi: 10.3390/ijms26136302.

57. Thirawatananond P, Brown ME, Sachs LK, Arnoletti JM, Yeh WI, Posgai AL, et al. Treg- Specific Cd226 Deletion Reduces Diabetes Incidence in Nod Mice by Improving Regulatory T-Cell Stability. Diabetes (2023) 72(11):1629–40. doi: 10.2337/db23-0307.

58. Ryoo HD. The Integrated Stress Response in Metabolic Adaptation. J Biol Chem (2024) 300(4):107151. Epub 20240309. doi: 10.1016/j.jbc.2024.107151.

59. Parashar S, Oliaeimotlagh M, Roy P, Lyu Q, Bellapu A, Fomin M, et al. Er Stress Induced Mitochondrial Dysfunction Drives Treg Instability in Coronary Artery Disease. EMBO Mol Med (2025) 17(12):3250–74. Epub 20251021. doi: 10.1038/s44321-025-00322-3.

60. Marciniak SJ, Yun CY, Oyadomari S, Novoa I, Zhang Y, Jungreis R, et al. Chop Induces Death by Promoting Protein Synthesis and Oxidation in the Stressed Endoplasmic Reticulum. Genes Dev (2004) 18(24):3066–77. doi: 10.1101/gad.1250704.

61. Schiavinato J, Haddad R, Saldanha-Araujo F, Baiochi J, Araujo AG, Santos Scheucher P, et al. Tgf-Beta/Atra-Induced Tregs Express a Selected Set of Micrornas Involved in the Repression of Transcripts Related to Th17 Differentiation. Sci Rep (2017) 7(1):3627. Epub 20170615. doi: 10.1038/s41598-017-03456-8.

62. Lee DY, Lee KS, Lee HJ, Kim DH, Noh YH, Yu K, et al. Activation of Perk Signaling Attenuates Abeta-Mediated Er Stress. PLoS One (2010) 5(5):e10489. Epub 20100505. doi: 10.1371/journal.pone.0010489.

63. Vieira FG, Tassinari VR, Kidd JD, Moreno A, Thompson K, Perrin S, et al. Perk Modulation, with Gsk2606414, Sephin1 or Salubrinal, Failed to Produce Therapeutic Benefits in the Sod1g93a Mouse Model of Als. PLoS One (2024) 19(2):e0292190. Epub 20240215. doi: 10.1371/journal.pone.0292190.

64. Krishnamoorthy J, Rajesh K, Mirzajani F, Kesoglidou P, Papadakis AI, Koromilas AE. Evidence for Eif2alpha Phosphorylation-Independent Effects of Gsk2656157, a Novel Catalytic Inhibitor of Perk with Clinical Implications. Cell Cycle (2014) 13(5):801–6. Epub 20140108. doi: 10.4161/cc.27726.

65. Lin S, Wu J, Lian G, Wu W, Chen W, Chen A, et al. Protective Effects of Salubrinal against H(2)O(2)-Induced Muscle Wasting Via Eif2alpha/Atf4 Signaling Pathway. Front Pharmacol (2025) 16:1607606. Epub 20250627. doi: 10.3389/fphar.2025.1607606.

66. Luhr M, Torgersen ML, Szalai P, Hashim A, Brech A, Staerk J, et al. The Kinase Perk and the Transcription Factor Atf4 Play Distinct and Essential Roles in Autophagy Resulting from Tunicamycin-Induced Er Stress. J Biol Chem (2019) 294(20):8197–217. Epub 20190329. doi: 10.1074/jbc.RA118.002829.

67. Teske BF, Wek SA, Bunpo P, Cundiff JK, McClintick JN, Anthony TG, et al. The Eif2 Kinase Perk and the Integrated Stress Response Facilitate Activation of Atf6 During Endoplasmic Reticulum Stress. Mol Biol Cell (2011) 22(22):4390–405. Epub 20110914. doi: 10.1091/mbc.E11-06-0510.

68. Amodio G, Moltedo O, Fasano D, Zerillo L, Oliveti M, Di Pietro P, et al. Perk-Mediated Unfolded Protein Response Activation and Oxidative Stress in Park20 Fibroblasts. Front Neurosci (2019) 13:673. Epub 20190627. doi: 10.3389/fnins.2019.00673.

