## supplemental fig 1 for "Pharmacologic and Genetic Modulation of PERK-associated Stress Signaling Enhances Regulatory T Cell Responses and Attenuates Intestinal Inflammation"

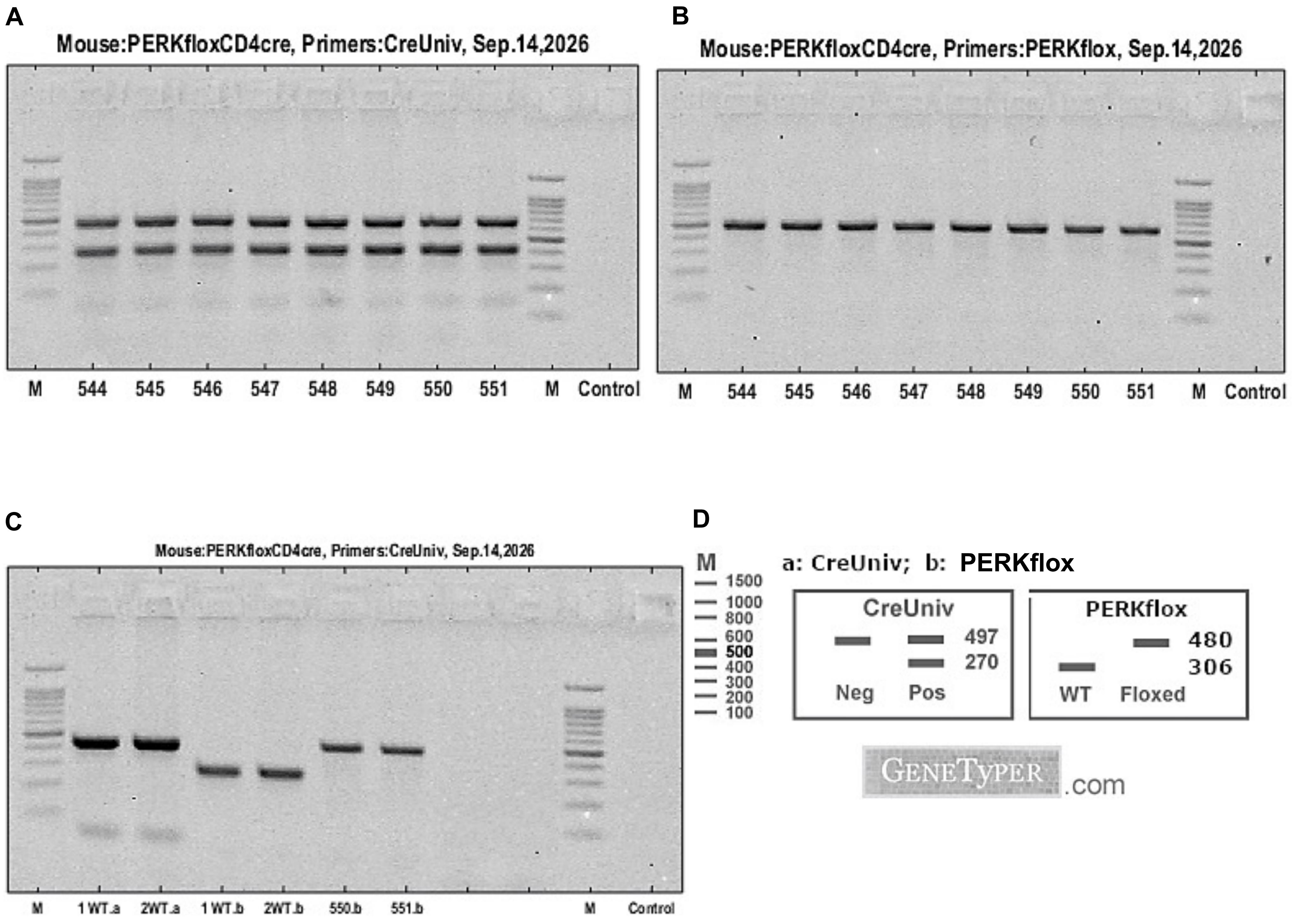
A
Mouse:PERKfloxCD4cre, Primers:CreUniv, Sep.14,2026
M 544 545 546 547 548 549 550 551 M Control
B
Mouse:PERKfloxCD4cre, Primers:PERKflox, Sep.14,2026
M 544 545 546 547 548 549 550 551 M Control
C
Mouse:PERKfloxCD4cre, Primers:CreUniv, Sep.14,2026
M 1WT.a 2WT.a 1WT.b 2WT.b 550.b 551.b M Control
D
M
1500
1000
800
600
500
400
300
200
100
a: CreUniv; b: PERKflox
CreUniv
497
270
Neg Pos
PERKflox
480
306
WT Floxed
GENETYPER.com
