## supplemental fig 2 for "Pharmacologic and Genetic Modulation of PERK-associated Stress Signaling Enhances Regulatory T Cell Responses and Attenuates Intestinal Inflammation"

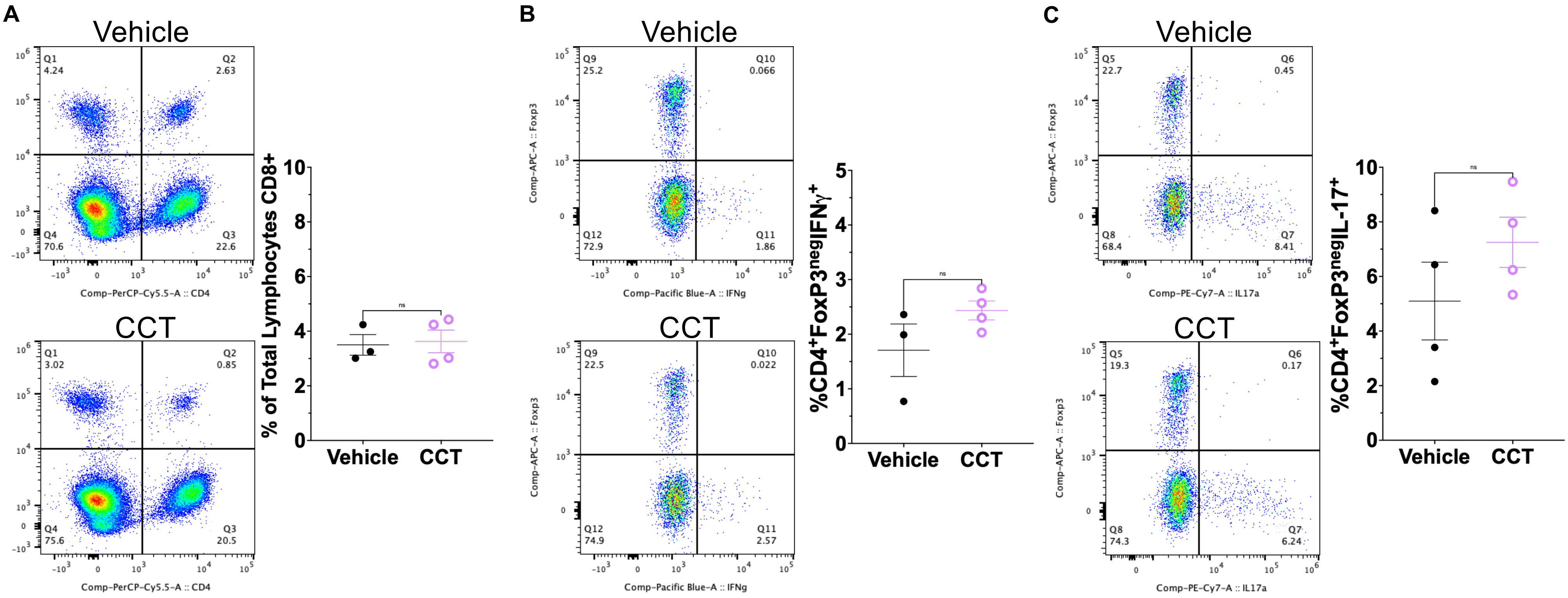
A
Vehicle
Q1
4.24
Q2
2.63
Q4
70.6
Q3
22.6
Comp-APC-A :: CD8
Comp-PerCP-Cy5.5-A :: CD4
CCT
Q1
3.02
Q2
0.85
Q4
75.6
Q3
20.5
% of Total Lymphocytes CD8+
ns
Vehicle
CCT
B
Vehicle
Q9
25.2
Q10
0.066
Q12
72.9
Q11
1.86
Comp-APC-A :: Foxp3
Comp-Pacific Blue-A :: IFNg
CCT
Q9
22.5
Q10
0.022
Q12
74.9
Q11
2.57
%CD4+FoxP3negIFNγ+
ns
Vehicle
CCT
C
Vehicle
Q5
22.7
Q6
0.45
Q8
68.4
Q7
8.41
Comp-APC-A :: Foxp3
Comp-PE-Cy7-A :: IL17a
CCT
Q5
19.3
Q6
0.17
Q8
74.3
Q7
6.24
%CD4+FoxP3negIL-17+
ns
Vehicle
CCT
