## supplemental fig 3 for "Pharmacologic and Genetic Modulation of PERK-associated Stress Signaling Enhances Regulatory T Cell Responses and Attenuates Intestinal Inflammation"

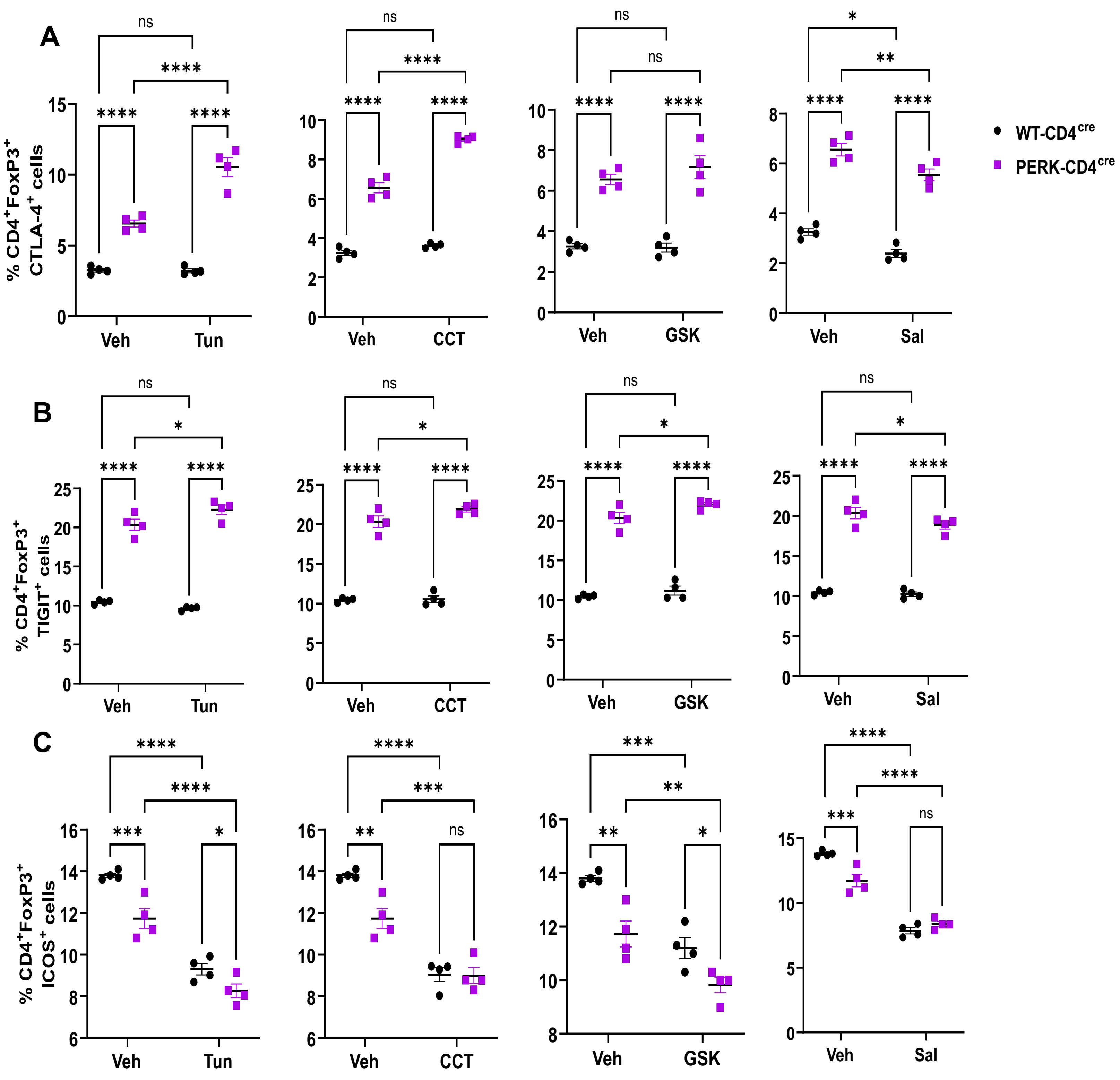
A
B
C
% CD4+FoxP3+ CTLA-4+ cells
% CD4+FoxP3+ TIGIT+ cells
% CD4+FoxP3+ ICOS+ cells
Veh
Tun
CCT
GSK
Sal
ns
*
**
***
****
WT-CD4cre
PERK-CD4cre
