## supplemental fig 4 for "Pharmacologic and Genetic Modulation of PERK-associated Stress Signaling Enhances Regulatory T Cell Responses and Attenuates Intestinal Inflammation"

| Experiment / figure | Biological system / input | Conditions / treatment | Exposure / timepoint | Primary readouts | n |
|---|---|---|---|---|---|
| Global UPR induction and iTreg conversion (Fig. 1) | WT $CD4^+CD25^{neg}$ T cells cultured under TGFβ1/IL-2/TCR iTreg conditions | Veh vs Tun (0.15 ng/mL), low glucose (0.5 g/L), or Thaps (50 nM) | UPR modulators for 4 h during conversion; total iTreg conversion 72 h | Flow cytometry for $\%CD4^+FoxP3^+$ Tregs | 3 biological replicates |
| Bulk RNA-seq (Fig. 2) | WT $CD4^+$ T cells from spleen and MLN | Veh vs Tun | 4 h at $37^0$ C | PCA, differential expression, heatmap, pathway grouping, GSVA correlation | 3 independent biological replicates |
| PERK modulation and Treg phenotyping (Fig. 3) | WT $CD4^+CD25^{neg}$ T cells in iTreg conversion assay | Veh, Tun, CCT (800 nM), GSK (1 uM), Sal (1 ug/mL) | Drug exposure during conversion; phenotyping at 72 h | Flow cytometry for $\%FoxP3^+$ Tregs and Neuropilin-1, ICOS, CTLA-4, TIGIT | 6 experimental replicates from 2 independent experiments |
| Immunoblot analysis of PERK-associated proteins (Fig. 4) | Converted Treg samples from iTreg assay | Veh, Tun, CCT, GSK, Sal | Protein harvested at assay endpoint | Immunoblot for p-PERK, p-eIF2α, FoxP3, IL-10, ATF4, CHOP; densitometry normalized to GAPDH | 3 biological replicates |
| Treg suppression assay (Fig. 5) | Converted Tregs co-cultured with CellTrace-labeled $CD4^+CD25^{neg}$ effector T cells and irradiated APCs | Veh, CCT, GSK, or Sal in Treg preparation; Treg:Teff ratios 1:1 to 1:64 | 72 h co-culture | Effector T-cell proliferation / suppression by CellTrace dilution | 3 experimental replicates |
| Single-cell RNA-seq of ileal LP T cells (Fig. 6) | Live ileal lamina propria cells from $TNF^{\Delta ARE+/-}$ mice; computational CD4 subset and Treg subset | Unmanipulated in vivo leukocytes; Tregs stratified as PERK_low vs PERK_high by module-score tertiles | Single endpoint at tissue harvest | UMAP, PERK module score, ISR gene violins, and DE heatmap between PERK_low and PERK_high Tregs | Each cells |
| In vivo PERK activation in $TNF^{\Delta ARE+/-}$ mice (Fig. 7) | Male $TNF^{\Delta ARE+/-}$ mice, 16-18 weeks old | Veh (DMSO in 1x PBS) vs CCT 4 mg/kg by daily i.p. injection | 14 consecutive days; harvest on day 14 | LP flow cytometry for $CD4^+FoxP3^+$ Tregs and $IL\text{-}10^+$ $CD4^+FoxP3^+$ Tregs; blinded ileal H&E histology | 3 Veh and 4 CCT biological replicates |
